## supplementary Information for "Functional Regulation of Aquaporin Dynamics by Lipid Bilayer Composition"

### Table of Contents


| | | |
|---|---|---|
| **Table S1** | Final Markov state model (MSM) parameters and total simulation time for each SoPIP2;1 lipid bilayer system | S-5 |
| **Table S2** | SoPIP2:POPC adaptive sampling summary | S-6 |
| **Table S3** | SoPIP2:POPE adaptive sampling summary | S-7 |
| **Table S4** | SoPIP2:POPG adaptive sampling summary | S-8 |
| **Table S5** | SoPIP2:PLPC adaptive sampling summary | S-9 |
| **Table S6** | SoPIP2:PLPE adaptive sampling summary | S-10 |
| **Table S7** | SoPIP2:PLPG adaptive sampling summary | S-11 |
| **Table S8** | SoPIP2:LLPC adaptive sampling summary | S-12 |
| **Table S9** | SoPIP2:LLPE adaptive sampling summary | S-13 |
| **Table S10** | SoPIP2:LLPG adaptive sampling summary | S-14 |
| **Table S11** | SoPIP2:complex adaptive sampling summary | S-15 |
| **Figure S1** | SoPIP2;1 topology | S-16 |
| **Figure S2** | Distance features and their correlation to tICA components generated by RRCS and spectral oASIS workflow | S-17 |
| **Figure S3** | Implied timescale plots for the SoPIP2:bilayer MSMs capturing the transitions for loop D conformational change. | S-18 |
| **Figure S4** | Raw counts versus MSM population for each microstate cluster of SoPIP2:bilayer conformational dynamics | S-19 |
| **Figure S5** | Chapman–Kolmogorov test for the MSM of SoPIP2:POPC conformational dynamics | S-20 |
| **Figure S6** | Chapman–Kolmogorov test for the MSM of SoPIP2:POPE conformational dynamics | S-21 |
| **Figure S7** | Chapman–Kolmogorov test for the MSM of SoPIP2:POPG conformational dynamics | S-22 |

**Figure S8** Chapman–Kolmogorov test for the MSM of SoPIP2:PLPC conformational dynamics S-23

**Figure S9** Chapman–Kolmogorov test for the MSM of SoPIP2:PLPE conformational dynamics S-24

**Figure S10** Chapman–Kolmogorov test for the MSM of SoPIP2:PLPG conformational dynamics S-25

**Figure S11** Chapman–Kolmogorov test for the MSM of SoPIP2:LLPC conformational dynamics S-26

**Figure S12** Chapman–Kolmogorov test for the MSM of SoPIP2:LLPE conformational dynamics S-27

**Figure S13** Chapman–Kolmogorov test for the MSM of SoPIP2:LLPG conformational dynamics S-28

**Figure S14** Chapman–Kolmogorov test for the MSM of SoPIP2:complex conformational dynamics S-29

**Figure S15** Free energy error from the 200-sample bootstrapping protocol for each SoPIP2:bilayer system S-30

**Figure S16** Selected continuous trajectories for the water transport analysis based on tICA macrostate identification S-31

**Figure S17** Relationship between number of waters transported and the rate of transport S-32

**Figure S18** Pore cavity structure with respect to loop D conformation throughout different types of example transport cases S-33

**Figure S19** Residence time of water molecules at each slice of the SoPIP2;1 pore S-34

**Figure S20** Dissecting influence of lipid headgroups or acyl chains on SoPIP2;1 protein function S-35

**Figure S21** Number of waters imported versus average lipid order parameter for all selected trajectories belonging to each SoPIP2:bilayer macrostate S-36

**Figure S22** Number of waters imported versus the average lipid order parameter of the annular shell lipids of the open states S-37

**Figure S23** Correlation between the thickness and order parameter of the lipids for each homogeneous SoPIP2:bilayer macrostate S-38

**Figure S24** Number of waters imported versus thickness for each SoPIP2:bilayer macrostate S-39

**Table S1. Final Markov state model (MSM) parameters and total simulation time for each SoPIP2;1 lipid bilayer system**

| Bilayer | Features | tIC dimensions | Clusters | MSM lag time (ns) | Total simulation time ($\mu$s) |
|---|---|---|---|---|---|
| POPC | 40 | 5 | 400 | 4 | 19.23 |
| POPE | 25 | 12 | 300 | 4 | 31.59 |
| POPG | 30 | 7 | 500 | 4 | 20.00 |
| PLPC | 30 | 7 | 700 | 6 | 54.71 |
| PLPE | 25 | 8 | 500 | 6 | 29.50 |
| PLPG | 40 | 4 | 500 | 4 | 26.30 |
| LLPC | 40 | 4 | 300 | 4 | 19.77 |
| LLPE | 25 | 7 | 400 | 6 | 49.50 |
| LLPG | 40 | 4 | 600 | 8 | 23.47 |
| Complex | 20 | 6 | 400 | 6 | 41.62 |

**Table S2. SoPIP2:POPC adaptive sampling summary.** The first round of simulation, "Round 1", consists of a single 1 μs trajectory originated from each of the crystal structure embeddings. This single trajectory took a few "rounds" to complete due to wallclock time limitations. The first "true" round of adaptive sampling is labelled as "Round 4". Each trajectory within a round of sampling outside of "Round 1" is set to be 100 ns in length.

| POPC | Parallel Simulations | | Trajectory Length (ns) | | Round Simulation Time (ns) | | Aggregate Time (μs) |
|---|---|---|---|---|---|---|---|
| Round | 2b5f (open) | 1z98 (closed) | 2b5f (open) | 1z98 (closed) | 2b5f (open) | 1z98 (closed) | |
| ***1*** | 1 | 1 | 1000 | 1000 | 1028.95 | 1200.00 | 2.23 |
| ***4*** | 2 | 8 | 100 | 100 | 200.00 | 800.00 | 1.00 |
| ***5*** | 3 | 7 | 100 | 100 | 300.00 | 700.00 | 1.00 |
| ***6*** | 10 | 10 | 100 | 100 | 1000.00 | 1000.00 | 2.00 |
| ***7*** | 10 | 10 | 100 | 100 | 1000.00 | 1000.00 | 2.00 |
| ***8*** | 10 | 10 | 100 | 100 | 1000.00 | 1000.00 | 2.00 |
| ***9*** | 5 | 5 | 100 | 100 | 500.00 | 500.00 | 1.00 |
| ***10*** | 5 | 5 | 100 | 100 | 500.00 | 500.00 | 1.00 |
| ***11*** | 10 | 10 | 100 | 100 | 1000.00 | 1000.00 | 2.00 |
| ***12*** | 0 | 10 | 100 | 100 | 0.00 | 1000.00 | 1.00 |
| ***13*** | 10 | 10 | 100 | 100 | 1000.00 | 1000.00 | 2.00 |
| ***14*** | 10 | 10 | 100 | 100 | 1000.00 | 1000.00 | 2.00 |
| **Total Simulation Time:** *19.23 μs* | | | | | | | |

**Table S3. SoPIP2:POPE adaptive sampling summary.** The first round of simulation, "Round 1", consists of a single 1 μs trajectory originated from each of the crystal structure embeddings. This single trajectory took a few "rounds" to complete due to wallclock time limitations. The first "true" round of adaptive sampling is labelled as "Round 4". Each trajectory within a round of sampling outside of "Round 1" is set to be 100 ns in length.

| POPE Round | Parallel Simulations | | Trajectory Length (ns) | | Round Simulation Time (ns) | | Aggregate Time (μs) |
|---|---|---|---|---|---|---|---|
| | 2b5f (open) | 1z98 (closed) | 2b5f (open) | 1z98 (closed) | 2b5f (open) | 1z98 (closed) | |
| ***1*** | 1 | 1 | 1000 | 1000 | 1000.00 | 1000.00 | 2.00 |
| ***4*** | 8 | 2 | 100 | 100 | 800.00 | 169.25 | 0.97 |
| ***5*** | 8 | 3 | 100 | 100 | 700.00 | 301.06 | 1.00 |
| ***6*** | 10 | 10 | 100 | 100 | 1000.00 | 1000.00 | 2.00 |
| ***7*** | 10 | 10 | 100 | 100 | 1000.00 | 921.26 | 1.92 |
| ***8*** | 10 | 10 | 100 | 100 | 1000.00 | 1000.00 | 2.00 |
| ***9*** | 10 | 10 | 100 | 100 | 1000.00 | 1000.00 | 2.00 |
| ***10*** | 10 | 10 | 100 | 100 | 1000.00 | 1000.00 | 2.00 |
| ***11*** | 10 | 10 | 100 | 100 | 1000.00 | 1000.00 | 2.00 |
| ***12*** | 10 | 10 | 100 | 100 | 1000.00 | 1000.00 | 2.00 |
| ***13*** | 7 | 10 | 100 | 100 | 700.00 | 1000.00 | 1.70 |
| ***14*** | 10 | 10 | 100 | 100 | 1000.00 | 1000.00 | 2.00 |
| ***15*** | 10 | 10 | 100 | 100 | 1000.00 | 1000.00 | 2.00 |
| ***16*** | 10 | 10 | 100 | 100 | 1000.00 | 1000.00 | 2.00 |
| ***17*** | 0 | 10 | 100 | 100 | 0.00 | 1000.00 | 1.00 |
| ***18*** | 0 | 10 | 100 | 100 | 0.00 | 1000.00 | 1.00 |
| ***19*** | 10 | 10 | 100 | 100 | 1000.00 | 1000.00 | 2.00 |
| ***20*** | 10 | 0 | 100 | 100 | 1000.00 | 0.00 | 1.00 |
| ***21*** | 10 | 0 | 100 | 100 | 1000.00 | 0.00 | 1.00 |
| **Total Simulation Time:** *31.59 μs* | | | | | | | |

**Table S4. SoPIP2:POPG adaptive sampling summary.** The first round of simulation, "Round 1", consists of a single 1 μs trajectory originated from each of the crystal structure embeddings. This single trajectory took a few "rounds" to complete due to wallclock time limitations. The first "true" round of adaptive sampling is labelled as "Round 4". Each trajectory within a round of sampling outside of "Round 1" is set to be 100 ns in length.

| POPG Round | Parallel Simulations | | Trajectory Length (ns) | | Round Simulation Time (ns) | | Aggregate Time (μs) |
|---|---|---|---|---|---|---|---|
| | 2b5f (open) | 1z98 (closed) | 2b5f (open) | 1z98 (closed) | 2b5f (open) | 1z98 (closed) | |
| ***1*** | 1 | 1 | 1000 | 1000 | 1000.00 | 1000.00 | 2.00 |
| ***4*** | 5 | 5 | 100 | 100 | 500.00 | 500.00 | 1.00 |
| ***5*** | 9 | 1 | 100 | 100 | 900.00 | 100.00 | 1.00 |
| ***6*** | 7 | 3 | 100 | 100 | 700.00 | 300.00 | 1.00 |
| ***7*** | 5 | 5 | 100 | 100 | 500.00 | 500.00 | 1.00 |
| ***8*** | 10 | 10 | 100 | 100 | 1000.00 | 1000.00 | 2.00 |
| ***9*** | 10 | 10 | 100 | 100 | 1000.00 | 1000.00 | 2.00 |
| ***10*** | 20 | 20 | 100 | 100 | 2000.00 | 2000.00 | 4.00 |
| ***11*** | 20 | 20 | 100 | 100 | 2000.00 | 2000.00 | 4.00 |
| ***12*** | 10 | 10 | 100 | 100 | 1000.00 | 1000.00 | 2.00 |
| **Total Simulation Time:** *20.00 μs* | | | | | | | |

**Table S5. SoPIP2:PLPC adaptive sampling summary.** The first round of simulation, "Round 1", consists of a single 1 μs trajectory originated from each of the crystal structure embeddings. This single trajectory took a few "rounds" to complete due to wallclock time limitations. The first "true" round of adaptive sampling is labelled as "Round 4". Each trajectory within a round of sampling outside of "Round 1" is set to be 100 ns in length.

| PLPC Round | Parallel Simulations | | Trajectory Length (ns) | | Round Simulation Time (ns) | | Aggregate Time (μs) |
|---|---|---|---|---|---|---|---|
| | 2b5f (open) | 1z98 (closed) | 2b5f (open) | 1z98 (closed) | 2b5f (open) | 1z98 (closed) | |
| ***1*** | 1 | 1 | 1000 | 1000 | 1000.00 | 1000.00 | 2.00 |
| ***4*** | 2 | 8 | 100 | 100 | 200.00 | 800.00 | 1.00 |
| ***5*** | 10 | 10 | 100 | 100 | 1000.00 | 1000.00 | 2.00 |
| ***6*** | 10 | 10 | 100 | 100 | 1000.00 | 1000.00 | 2.00 |
| ***7*** | 10 | 10 | 100 | 100 | 1000.00 | 1000.00 | 2.00 |
| ***8*** | 5 | 5 | 100 | 100 | 500.00 | 500.00 | 1.00 |
| ***9*** | 10 | 10 | 100 | 100 | 1000.00 | 1000.00 | 2.00 |
| ***10*** | 10 | 10 | 100 | 100 | 1000.00 | 1000.00 | 2.00 |
| ***11*** | 10 | 10 | 100 | 100 | 1000.00 | 1000.00 | 2.00 |
| ***12*** | 10 | 10 | 100 | 100 | 1000.00 | 1000.00 | 2.00 |
| ***13*** | 10 | 10 | 100 | 100 | 1000.00 | 1000.00 | 2.00 |
| ***14*** | 10 | 10 | 100 | 100 | 1000.00 | 1000.00 | 2.00 |
| ***15*** | 10 | 10 | 100 | 100 | 1000.00 | 1000.00 | 2.00 |
| ***16*** | 10 | 10 | 100 | 100 | 1000.00 | 1000.00 | 2.00 |
| ***17*** | 10 | 10 | 100 | 100 | 1000.00 | 1000.00 | 2.00 |
| ***18*** | 10 | 10 | 100 | 100 | 1000.00 | 1000.00 | 2.00 |
| ***19*** | 10 | 10 | 100 | 100 | 1000.00 | 1000.00 | 2.00 |
| ***20*** | 10 | 10 | 100 | 100 | 863.68 | 1000.00 | 1.86 |
| ***21*** | 10 | 10 | 100 | 100 | 996.34 | 952.20 | 1.95 |
| ***22*** | 10 | 10 | 100 | 100 | 1000.00 | 1000.00 | 2.00 |
| ***23*** | 10 | 10 | 100 | 100 | 1000.00 | 1000.00 | 2.00 |
| ***24*** | 10 | 10 | 100 | 100 | 1000.00 | 1000.00 | 2.00 |
| ***25*** | 10 | 0 | 100 | 100 | 1000.00 | 0.00 | 1.00 |
| ***26*** | 10 | 0 | 100 | 100 | 1000.00 | 0.00 | 1.00 |
| ***27*** | 10 | 0 | 100 | 100 | 1000.00 | 0.00 | 1.00 |
| ***28*** | 10 | 10 | 100 | 100 | 1000.00 | 1000.00 | 2.00 |
| ***29*** | 15 | 15 | 100 | 100 | 1500.00 | 1500.00 | 3.00 |
| ***30*** | 9 | 10 | 100 | 100 | 900.00 | 1000.00 | 1.90 |
| ***31*** | 15 | 15 | 100 | 100 | 1500.00 | 1500.00 | 3.00 |
| **Total Simulation Time:** *54.71 μs* | | | | | | | |

**Table S6. SoPIP2:PLPE adaptive sampling summary.** The first round of simulation, "Round 1", consists of a single 1 μs trajectory originated from each of the crystal structure embeddings. This single trajectory took a few "rounds" to complete due to wallclock time limitations. The first "true" round of adaptive sampling is labelled as "Round 4". Each trajectory within a round of sampling outside of "Round 1" is set to be 100 ns in length.

| **PLPE** | **Parallel Simulations** | | **Trajectory Length (ns)** | | **Round Simulation Time (ns)** | | **Aggregate Time (μs)** |
|---|---|---|---|---|---|---|---|
| **Round** | **2b5f (open)** | **1z98 (closed)** | **2b5f (open)** | **1z98 (closed)** | **2b5f (open)** | **1z98 (closed)** | |
| ***1*** | 1 | 1 | 1000 | 1000 | 1000.00 | 1000.00 | 2.00 |
| ***4*** | 10 | 10 | 100 | 100 | 1000.00 | 1000.00 | 2.00 |
| ***5*** | 10 | 0 | 100 | 100 | 1000.00 | 0.00 | 1.00 |
| ***6*** | 10 | 10 | 100 | 100 | 1000.00 | 1000.00 | 2.00 |
| ***7*** | 10 | 10 | 100 | 100 | 1000.00 | 1000.00 | 2.00 |
| ***8*** | 10 | 10 | 100 | 100 | 1000.00 | 1000.00 | 2.00 |
| ***9*** | 10 | 10 | 100 | 100 | 1000.00 | 1000.00 | 2.00 |
| ***10*** | 5 | 5 | 100 | 100 | 500.00 | 500.00 | 1.00 |
| ***11*** | 10 | 10 | 100 | 100 | 1000.00 | 1000.00 | 2.00 |
| ***12*** | 10 | 10 | 100 | 100 | 1000.00 | 1000.00 | 2.00 |
| ***13*** | 10 | 10 | 100 | 100 | 1000.00 | 1000.00 | 2.00 |
| ***14*** | 10 | 10 | 100 | 100 | 1000.00 | 1000.00 | 2.00 |
| ***15*** | 10 | 10 | 100 | 100 | 1000.00 | 1000.00 | 2.00 |
| ***16*** | 10 | 10 | 100 | 100 | 1000.00 | 1000.00 | 2.00 |
| ***17*** | 15 | 0 | 100 | 100 | 1500.00 | 0.00 | 1.50 |
| ***18*** | 20 | 0 | 100 | 100 | 2000.00 | 0.00 | 2.00 |
| **Total Simulation Time:** *29.50 μs* | | | | | | | |

**Table S7. SoPIP2:PLPG adaptive sampling summary.** The first round of simulation, "Round 1", consists of a single 1 μs trajectory originated from each of the crystal structure embeddings. This single trajectory took a few "rounds" to complete due to wallclock time limitations. The first "true" round of adaptive sampling is labelled as "Round 4". Each trajectory within a round of sampling outside of "Round 1" is set to be 100 ns in length.

| PLPG Round | Parallel Simulations | | Trajectory Length (ns) | | Round Simulation Time (ns) | | Aggregate Time (μs) |
|---|---|---|---|---|---|---|---|
| | 2b5f (open) | 1z98 (closed) | 2b5f (open) | 1z98 (closed) | 2b5f (open) | 1z98 (closed) | |
| ***1*** | 1 | 1 | 1000 | 1000 | 1000.00 | 900.00 | 1.90 |
| ***4*** | 8 | 2 | 100 | 100 | 800.00 | 200.00 | 1.00 |
| ***5*** | 10 | 10 | 100 | 100 | 1000.00 | 1000.00 | 2.00 |
| ***6*** | 10 | 0 | 100 | 100 | 1000.00 | 0.00 | 1.00 |
| ***7*** | 10 | 0 | 100 | 100 | 1000.00 | 0.00 | 1.00 |
| ***8*** | 10 | 0 | 100 | 100 | 1000.00 | 0.00 | 1.00 |
| ***9*** | 10 | 0 | 100 | 100 | 1000.00 | 0.00 | 1.00 |
| ***10*** | 7 | 3 | 100 | 100 | 700.00 | 300.00 | 1.00 |
| ***11*** | 20 | 0 | 100 | 100 | 2000.00 | 0.00 | 2.00 |
| ***12*** | 20 | 0 | 100 | 100 | 2000.00 | 0.00 | 2.00 |
| ***13*** | 10 | 0 | 100 | 100 | 1000.00 | 0.00 | 1.00 |
| ***14*** | 10 | 0 | 100 | 100 | 1000.00 | 0.00 | 1.00 |
| ***15*** | 0 | 20 | 100 | 100 | 0.00 | 2000.00 | 2.00 |
| ***16*** | 15 | 15 | 100 | 100 | 1500.00 | 1500.00 | 3.00 |
| ***17*** | 10 | 20 | 100 | 100 | 1000.00 | 2000.00 | 3.00 |
| ***18*** | 10 | 15 | 100 | 100 | 903.28 | 1500.00 | 2.40 |
| **Total Simulation Time:** *26.30 μs* | | | | | | | |

**Table S8. SoPIP2:LLPC adaptive sampling summary.** The first round of simulation, "Round 1", consists of a single 1 μs trajectory originated from each of the crystal structure embeddings. This single trajectory took a few "rounds" to complete due to wallclock time limitations. The first "true" round of adaptive sampling is labelled as "Round 4". Each trajectory within a round of sampling outside of "Round 1" is set to be 100 ns in length.

| LLPC Round | Parallel Simulations | | Trajectory Length (ns) | | Round Simulation Time (ns) | | Aggregate Time (μs) |
|---|---|---|---|---|---|---|---|
| | 2b5f (open) | 1z98 (closed) | 2b5f (open) | 1z98 (closed) | 2b5f (open) | 1z98 (closed) | |
| ***1*** | 1 | 1 | 1000 | 1000 | 914.33 | 1000.00 | 1.91 |
| ***4*** | 7 | 3 | 100 | 100 | 700.00 | 300.00 | 1.00 |
| ***5*** | 2 | 8 | 100 | 100 | 200.00 | 798.50 | 1.00 |
| ***6*** | 10 | 10 | 100 | 100 | 716.30 | 936.63 | 1.65 |
| ***7*** | 10 | 5 | 100 | 100 | 940.12 | 438.75 | 1.38 |
| ***8*** | 10 | 10 | 100 | 100 | 842.77 | 986.74 | 1.83 |
| ***9*** | 10 | 10 | 100 | 100 | 1000.00 | 1000.00 | 2.00 |
| ***10*** | 10 | 10 | 100 | 100 | 1000.00 | 1000.00 | 2.00 |
| ***11*** | 10 | 10 | 100 | 100 | 1000.00 | 1000.00 | 2.00 |
| ***12*** | 10 | 0 | 100 | 100 | 1000.00 | 0.00 | 1.00 |
| ***13*** | 10 | 10 | 100 | 100 | 1000.00 | 1000.00 | 2.00 |
| ***14*** | 10 | 10 | 100 | 100 | 1000.00 | 1000.00 | 2.00 |
| **Total Simulation Time:** *19.77 μs* | | | | | | | |

**Table S9. SoPIP2:LLPE adaptive sampling summary.** The first round of simulation, "Round 1", consists of a single 1 μs trajectory originated from each of the crystal structure embeddings. This single trajectory took a few "rounds" to complete due to wallclock time limitations. The first "true" round of adaptive sampling is labelled as "Round 4". Each trajectory within a round of sampling outside of "Round 1" is set to be 100 ns in length.

| LLPE Round | Parallel Simulations | | Trajectory Length (ns) | | Round Simulation Time (ns) | | Aggregate Time (μs) |
|---|---|---|---|---|---|---|---|
| | 2b5f (open) | 1z98 (closed) | 2b5f (open) | 1z98 (closed) | 2b5f (open) | 1z98 (closed) | |
| ***1*** | 1 | 1 | 1000 | 1000 | 1736.20 | 1000.00 | 2.74 |
| ***4*** | 10 | 10 | 100 | 100 | 1000.00 | 1000.00 | 2.00 |
| ***5*** | 10 | 10 | 100 | 100 | 1000.00 | 1000.00 | 2.00 |
| ***6*** | 10 | 10 | 100 | 100 | 1000.00 | 1000.00 | 2.00 |
| ***7*** | 10 | 10 | 100 | 100 | 1000.00 | 1000.00 | 2.00 |
| ***8*** | 4 | 5 | 100 | 100 | 400.00 | 500.00 | 0.90 |
| ***9*** | 8 | 8 | 100 | 100 | 800.00 | 800.00 | 1.60 |
| ***10*** | 13 | 5 | 100 | 100 | 1300.00 | 500.00 | 1.80 |
| ***11*** | 10 | 10 | 100 | 100 | 1000.00 | 1000.00 | 2.00 |
| ***12*** | 10 | 10 | 100 | 100 | 1000.00 | 1000.00 | 2.00 |
| ***13*** | 10 | 10 | 100 | 100 | 1000.00 | 1000.00 | 2.00 |
| ***14*** | 10 | 10 | 100 | 100 | 1000.00 | 1000.00 | 2.00 |
| ***15*** | 10 | 10 | 100 | 100 | 1000.00 | 1000.00 | 2.00 |
| ***16*** | 10 | 10 | 100 | 100 | 1000.00 | 1000.00 | 2.00 |
| ***17*** | 10 | 10 | 100 | 100 | 1000.00 | 1000.00 | 2.00 |
| ***18*** | 10 | 10 | 100 | 100 | 1000.00 | 1000.00 | 2.00 |
| ***19*** | 10 | 10 | 100 | 100 | 1000.00 | 1000.00 | 2.00 |
| ***20*** | 10 | 10 | 100 | 100 | 1000.00 | 1000.00 | 2.00 |
| ***21*** | 10 | 10 | 100 | 100 | 1000.00 | 1000.00 | 2.00 |
| ***22*** | 10 | 10 | 100 | 100 | 1000.00 | 1000.00 | 2.00 |
| ***23*** | 10 | 10 | 100 | 100 | 1000.00 | 963.42 | 1.96 |
| ***24*** | 10 | 10 | 100 | 100 | 1000.00 | 1000.00 | 2.00 |
| ***25*** | 10 | 10 | 100 | 100 | 1000.00 | 1000.00 | 2.00 |
| ***26*** | 0 | 10 | 100 | 100 | 0.00 | 1000.00 | 1.00 |
| ***27*** | 0 | 20 | 100 | 100 | 0.00 | 2000.00 | 2.00 |
| ***28*** | 15 | 0 | 100 | 100 | 1500.00 | 0.00 | 1.50 |
| **Total Simulation Time:** *49.50 μs* | | | | | | | |

**Table S10. SoPIP2:LLPG adaptive sampling summary.** The first round of simulation, "Round 1", consists of a single 1 μs trajectory originated from each of the crystal structure embeddings. This single trajectory took a few "rounds" to complete due to wallclock time limitations. The first "true" round of adaptive sampling is labelled as "Round 4". Each trajectory within a round of sampling outside of "Round 1" is set to be 100 ns in length.

| LLPG | Parallel Simulations | | Trajectory Length (ns) | | Round Simulation Time (ns) | | Aggregate Time (μs) |
|---|---|---|---|---|---|---|---|
| Round | 2b5f (open) | 1z98 (closed) | 2b5f (open) | 1z98 (closed) | 2b5f (open) | 1z98 (closed) | |
| ***1*** | 1 | 1 | 1000 | 1000 | 1000.00 | 1000.00 | 2.00 |
| ***4*** | 10 | 10 | 100 | 100 | 1000.00 | 1000.00 | 2.00 |
| ***5*** | 10 | 0 | 100 | 100 | 1000.00 | 0.00 | 1.00 |
| ***6*** | 10 | 10 | 100 | 100 | 1000.00 | 1000.00 | 2.00 |
| ***7*** | 10 | 10 | 100 | 100 | 1000.00 | 1000.00 | 2.00 |
| ***8*** | 10 | 10 | 100 | 100 | 1000.00 | 1000.00 | 2.00 |
| ***9*** | 10 | 10 | 100 | 100 | 1000.00 | 1000.00 | 2.00 |
| ***10*** | 5 | 5 | 100 | 100 | 500.00 | 500.00 | 1.00 |
| ***11*** | 5 | 6 | 100 | 100 | 500.00 | 600.00 | 1.10 |
| ***12*** | 10 | 0 | 100 | 100 | 1000.00 | 0.00 | 1.00 |
| ***13*** | 10 | 0 | 100 | 100 | 1000.00 | 0.00 | 1.00 |
| ***14*** | 10 | 10 | 100 | 100 | 1000.00 | 1000.00 | 2.00 |
| ***15*** | 0 | 10 | 100 | 100 | 0.00 | 1000.00 | 1.00 |
| ***16*** | 20 | 15 | 100 | 100 | 2000.00 | 1371.42 | 3.37 |
| **Total Simulation Time:** *23.47 μs* | | | | | | | |

**Table S11. SoPIP2:complex adaptive sampling summary.** The first round of simulation, “Round 1”, consists of a single 1 μs trajectory originated from each of the crystal structure embeddings. This single trajectory took a few “rounds” to complete due to wallclock time limitations. The first “true” round of adaptive sampling is labelled as “Round 4”. Each trajectory within a round of sampling outside of “Round 1” is set to be 100 ns in length.

| complex | Parallel Simulations | | Trajectory Length (ns) | | Round Simulation Time (ns) | | Aggregate Time (μs) |
|---|---|---|---|---|---|---|---|
| Round | 2b5f (open) | 1z98 (closed) | 2b5f (open) | 1z98 (closed) | 2b5f (open) | 1z98 (closed) | |
| ***1*** | 1 | 1 | 1000 | 1000 | 791.17 | 1200.00 | 1.99 |
| ***4*** | 6 | 4 | 100 | 100 | 600.00 | 400.00 | 1.00 |
| ***5*** | 10 | 10 | 60 | 100 | 664.30 | 1000.00 | 1.66 |
| ***6*** | 10 | 10 | 100 | 100 | 1000.00 | 1000.00 | 2.00 |
| ***7*** | 10 | 10 | 100 | 100 | 1000.00 | 1000.00 | 2.00 |
| ***8*** | 5 | 5 | 100 | 100 | 500.00 | 500.00 | 1.00 |
| ***9*** | 5 | 5 | 100 | 100 | 500.00 | 500.00 | 1.00 |
| ***10*** | 10 | 10 | 100 | 100 | 1000.00 | 1000.00 | 2.00 |
| ***11*** | 10 | 10 | 100 | 100 | 1000.00 | 1000.00 | 2.00 |
| ***12*** | 10 | 10 | 100 | 100 | 1000.00 | 1000.00 | 2.00 |
| ***13*** | 10 | 10 | 100 | 100 | 1000.00 | 1000.00 | 2.00 |
| ***14*** | 10 | 10 | 100 | 100 | 1000.00 | 1000.00 | 2.00 |
| ***15*** | 10 | 10 | 100 | 100 | 1000.00 | 1000.00 | 2.00 |
| ***16*** | 10 | 10 | 100 | 100 | 1000.00 | 1000.00 | 2.00 |
| ***17*** | 10 | 10 | 100 | 100 | 1000.00 | 1000.00 | 2.00 |
| ***18*** | 10 | 10 | 100 | 100 | 1000.00 | 1000.00 | 2.00 |
| ***19*** | 10 | 10 | 100 | 100 | 1000.00 | 1000.00 | 2.00 |
| ***20*** | 10 | 10 | 100 | 100 | 1000.00 | 1000.00 | 2.00 |
| ***21*** | 0 | 10 | 100 | 100 | 0.00 | 1000.00 | 1.00 |
| ***22*** | 0 | 10 | 100 | 100 | 0.00 | 961.39 | 0.96 |
| ***23*** | 10 | 10 | 100 | 100 | 1000.00 | 1000.00 | 2.00 |
| ***24*** | 15 | 15 | 100 | 100 | 1500.00 | 1500.00 | 3.00 |
| ***25*** | 10 | 10 | 100 | 100 | 1000.00 | 1000.00 | 2.00 |
| **Total Simulation Time:** *41.62 μs* | | | | | | | |

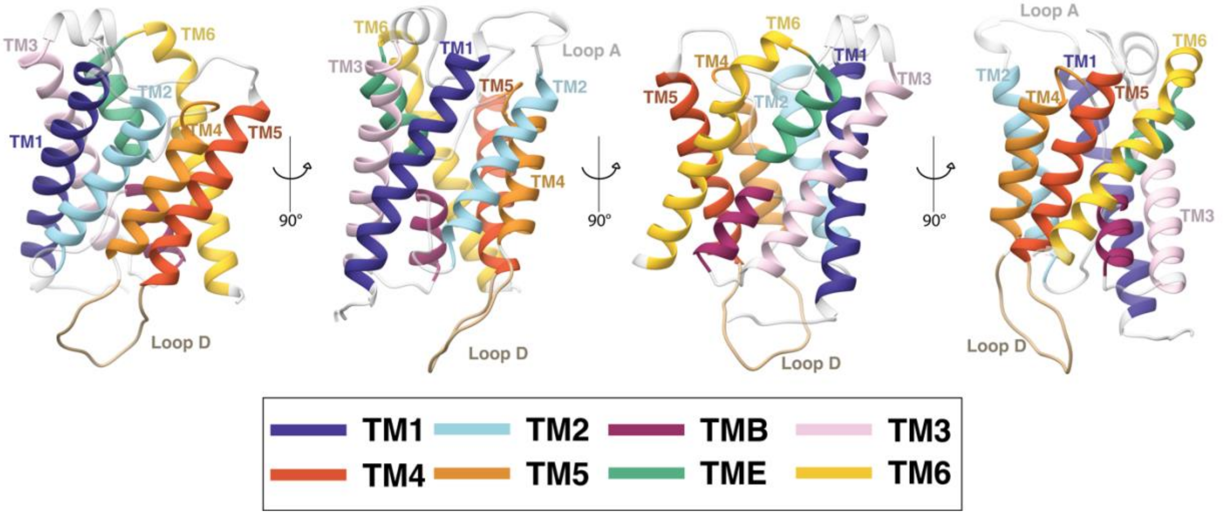


**Figure S1. SoPIP2;1 topology.** The helices and loop D are colored and labelled on the SoPIP2;1 open macrostate. Loop A is labelled when visible. TM indicates transmembrane domain according to the aquaporin topology. TMB and TME are two half helices in the lower leaflet and upper leaflet, respectively, of the lipid bilayer.

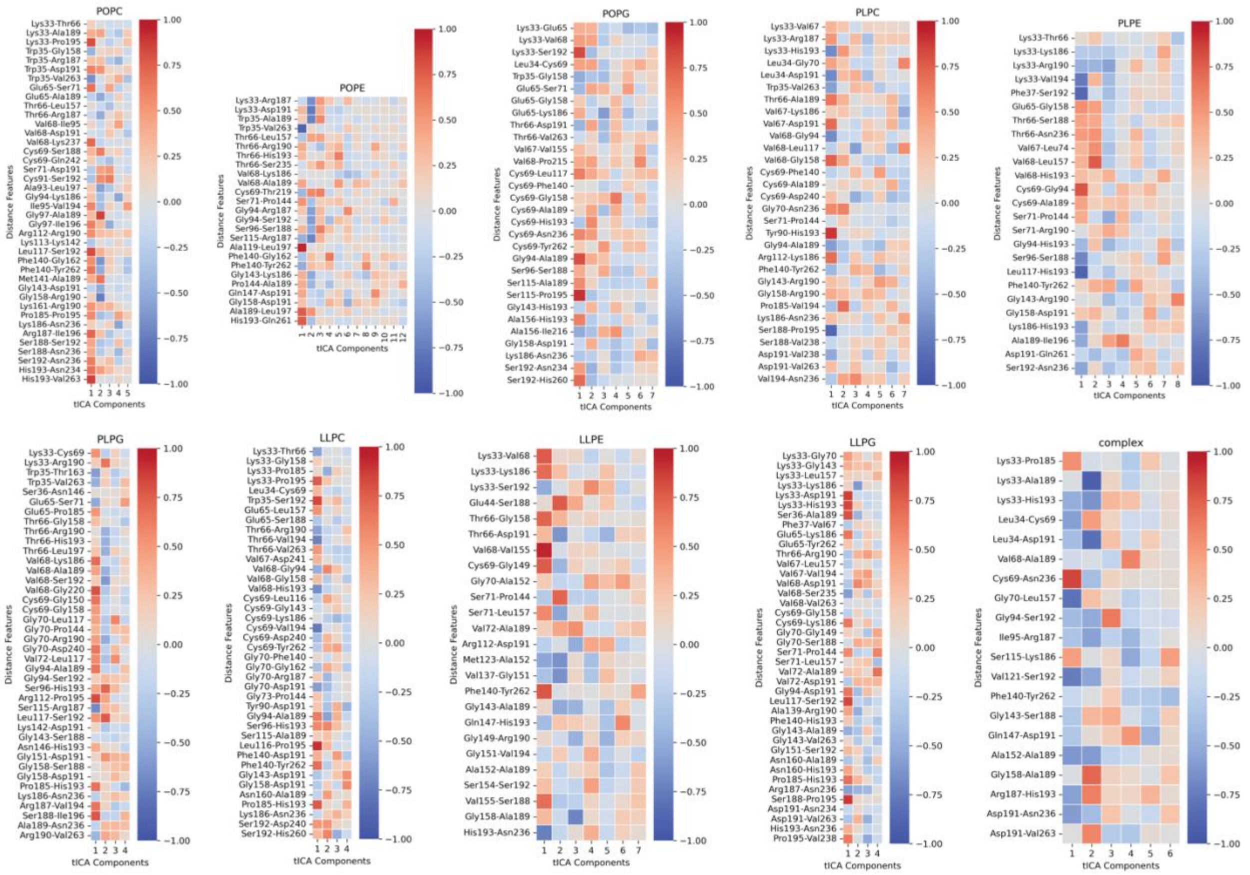


**Figure S2. Distance features and their correlation to tICA components generated by RRCS and spectral oASIS workflow.** All heatmaps represent correlations for features used in each of the final SoPIP2;1 MSMs.

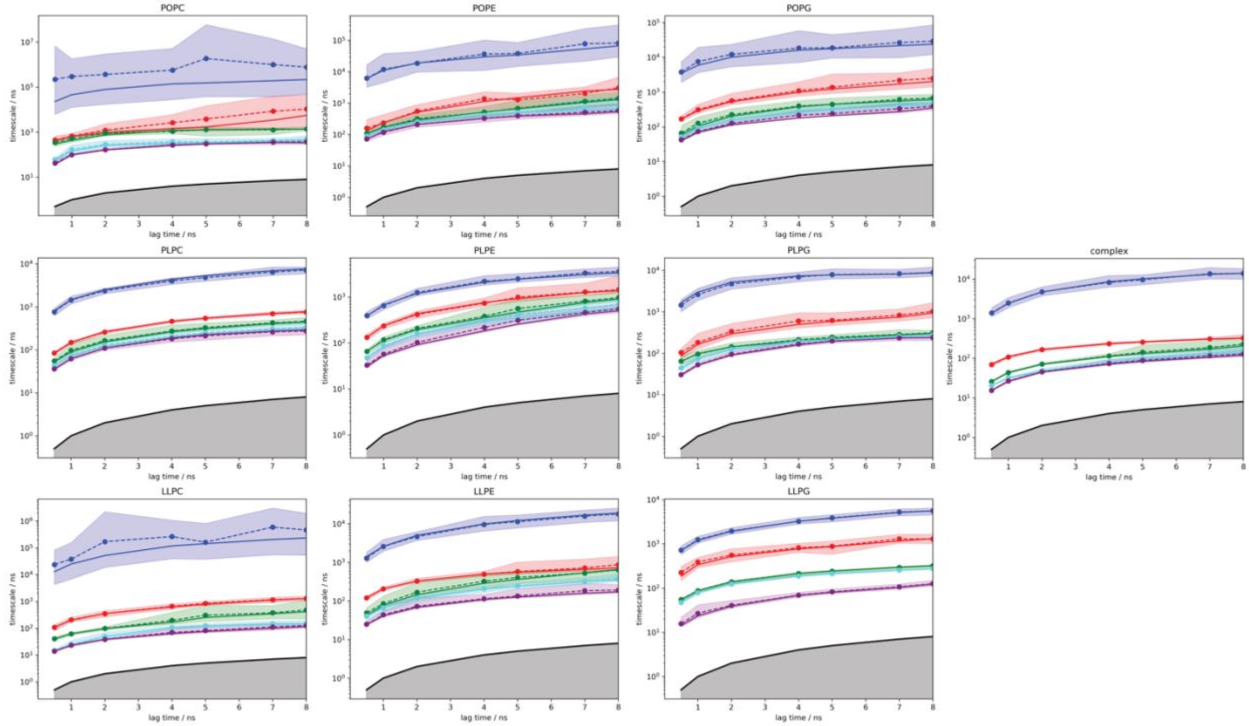


**Figure S3. Implied timescale plots for the SoPIP2:bilayer MSMs capturing the transitions for loop D conformational change.** Implied timescale plots were calculated with Bayesian error.

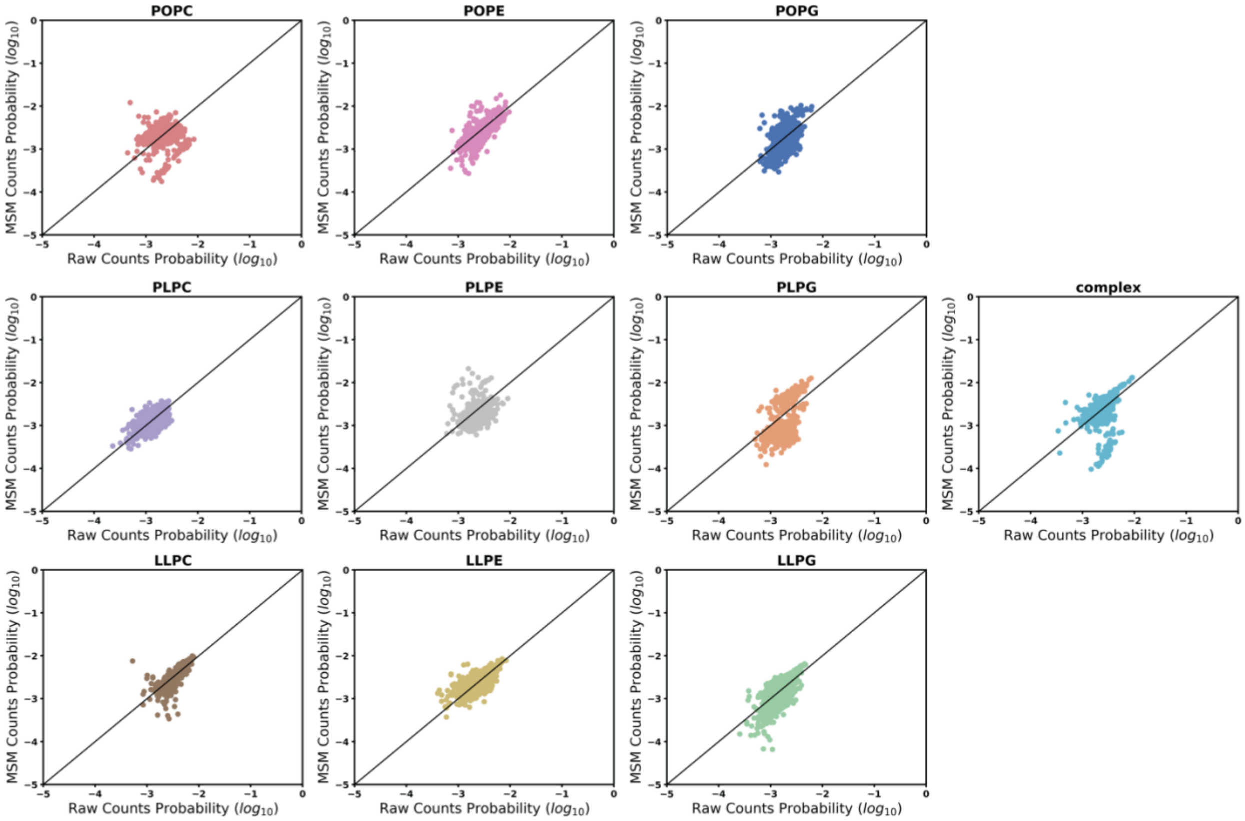


**Figure S4. Raw counts versus MSM population for each microstate cluster of SoPIP2:bilayer conformational dynamics.**

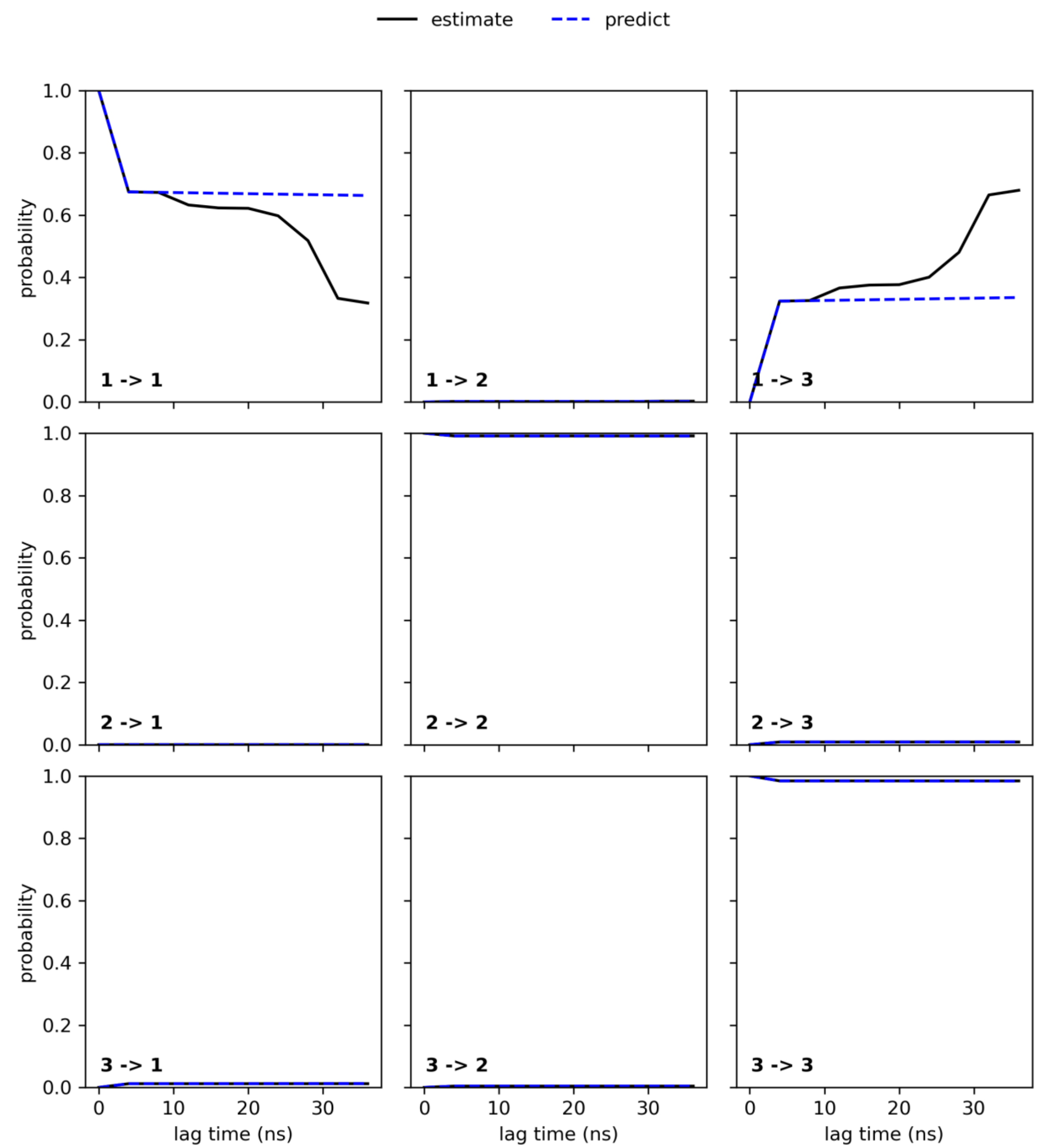


**Figure S5. Chapman–Kolmogorov test for the MSM of SoPIP2:POPC conformational dynamics.**

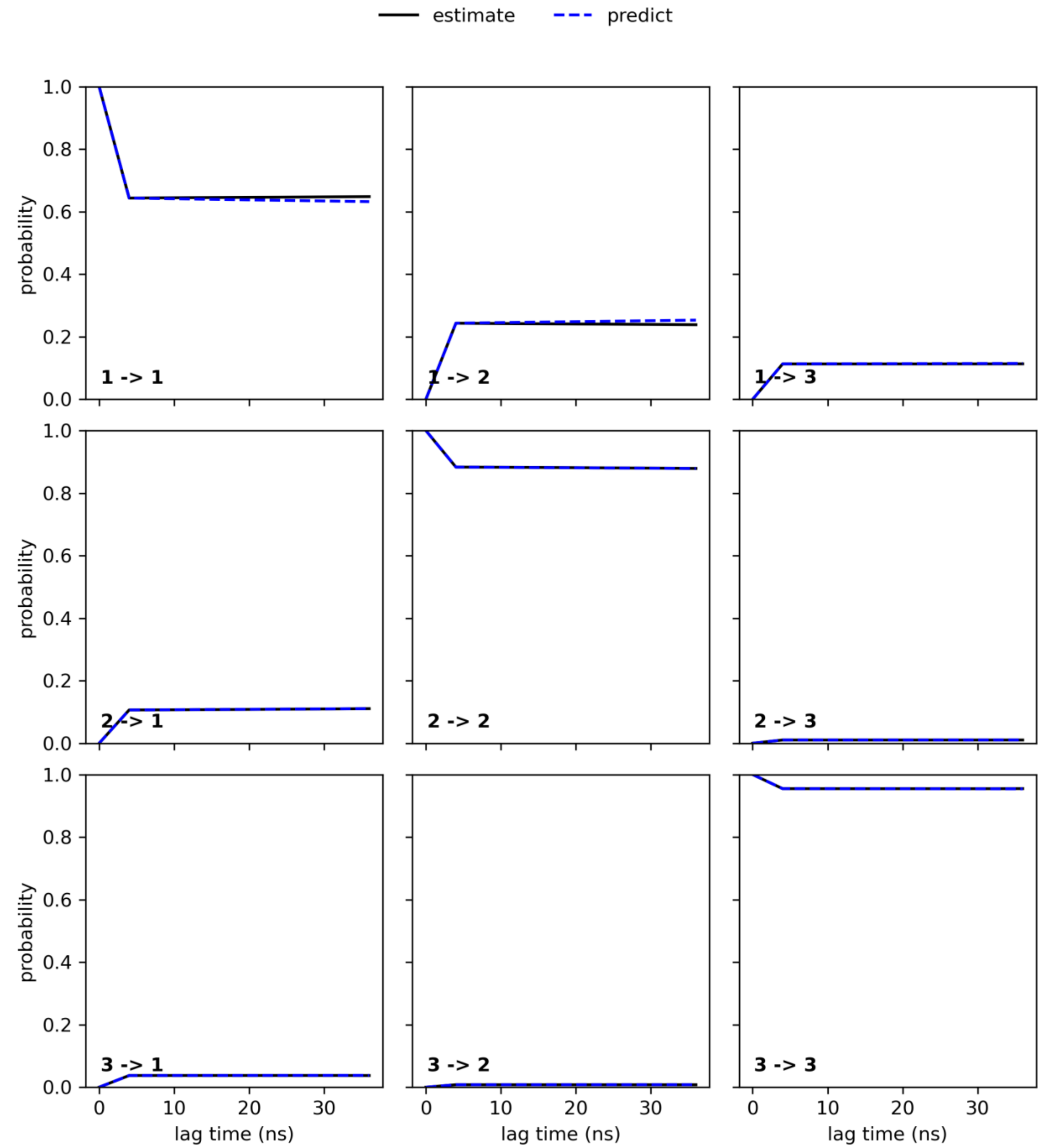


**Figure S6. Chapman–Kolmogorov test for the MSM of SoPIP2:POPE conformational dynamics.**

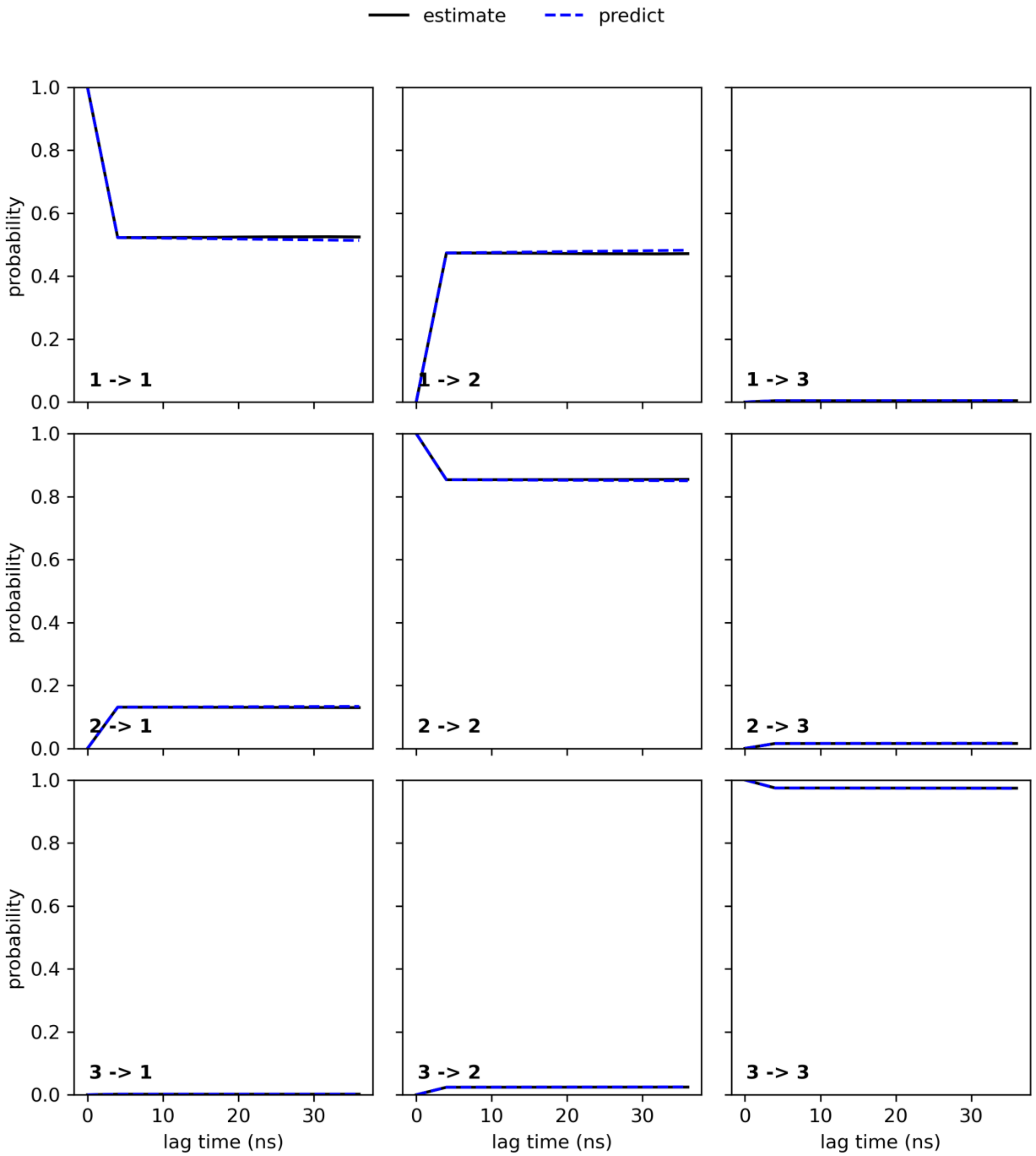


**Figure S7. Chapman–Kolmogorov test for the MSM of SoPIP2:POPG conformational dynamics.**

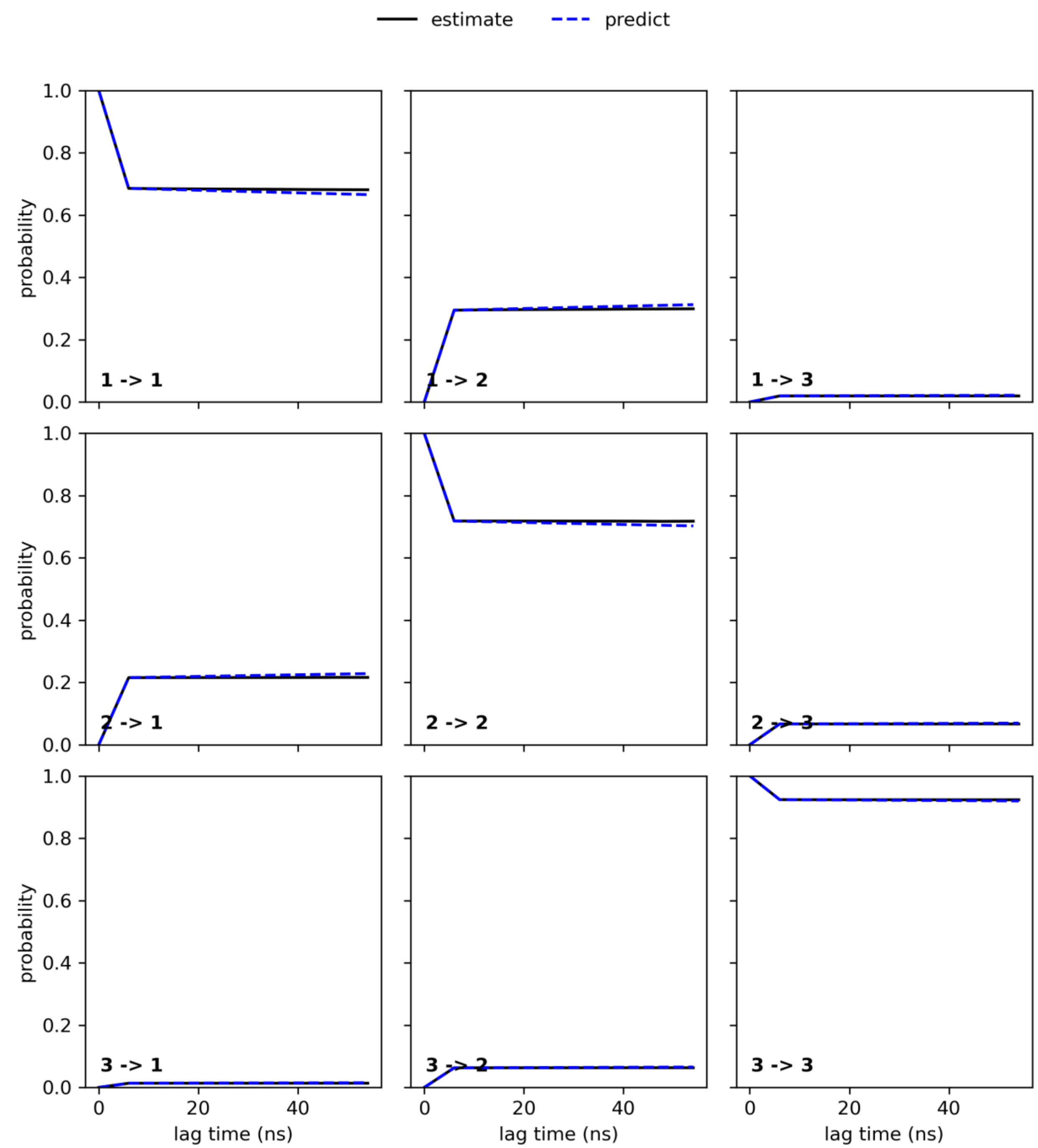


**Figure S8. Chapman–Kolmogorov test for the MSM of SoPIP2:PLPC conformational dynamics.**

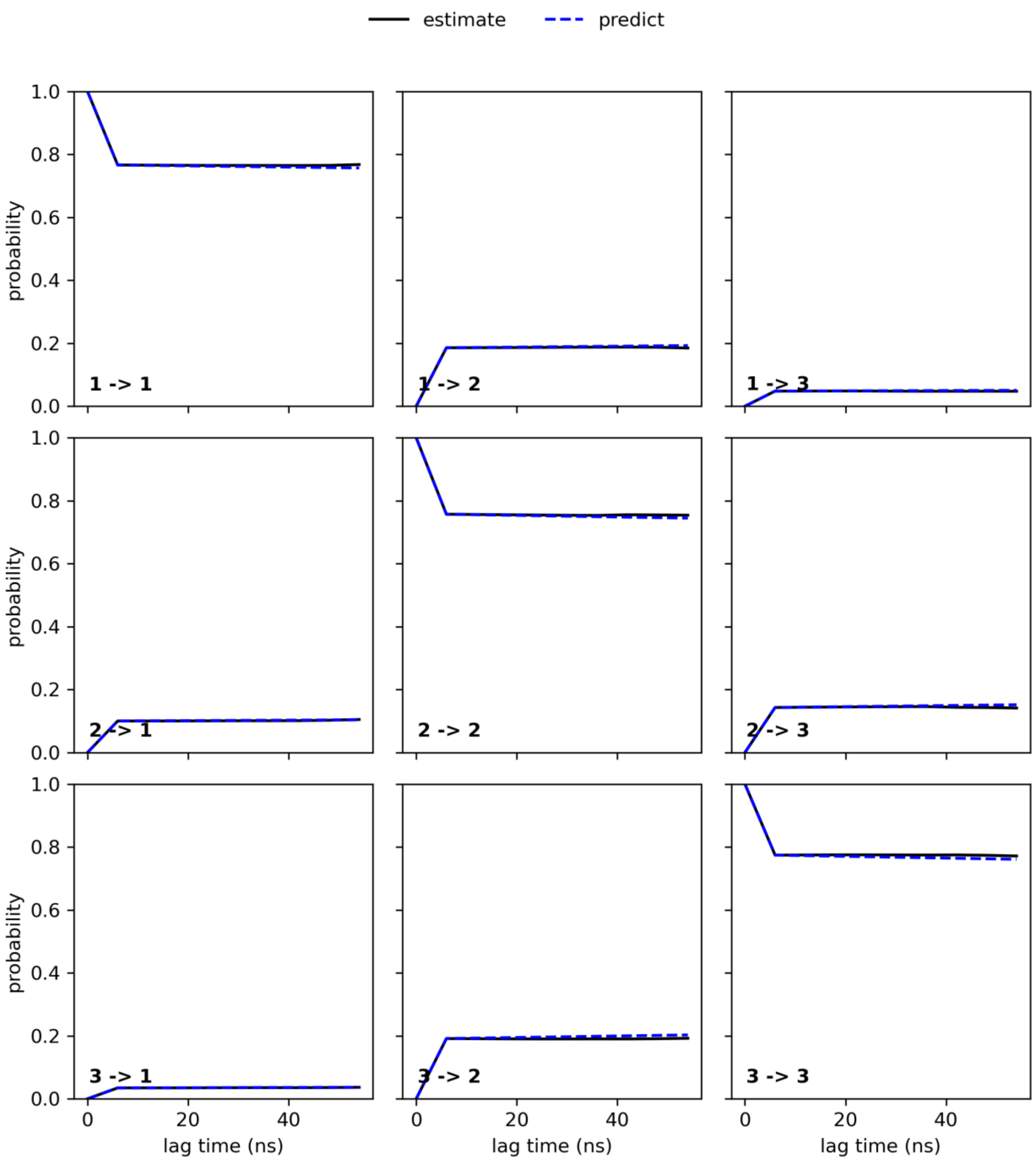


**Figure S9. Chapman–Kolmogorov test for the MSM of SoPIP2:PLPE conformational dynamics.**

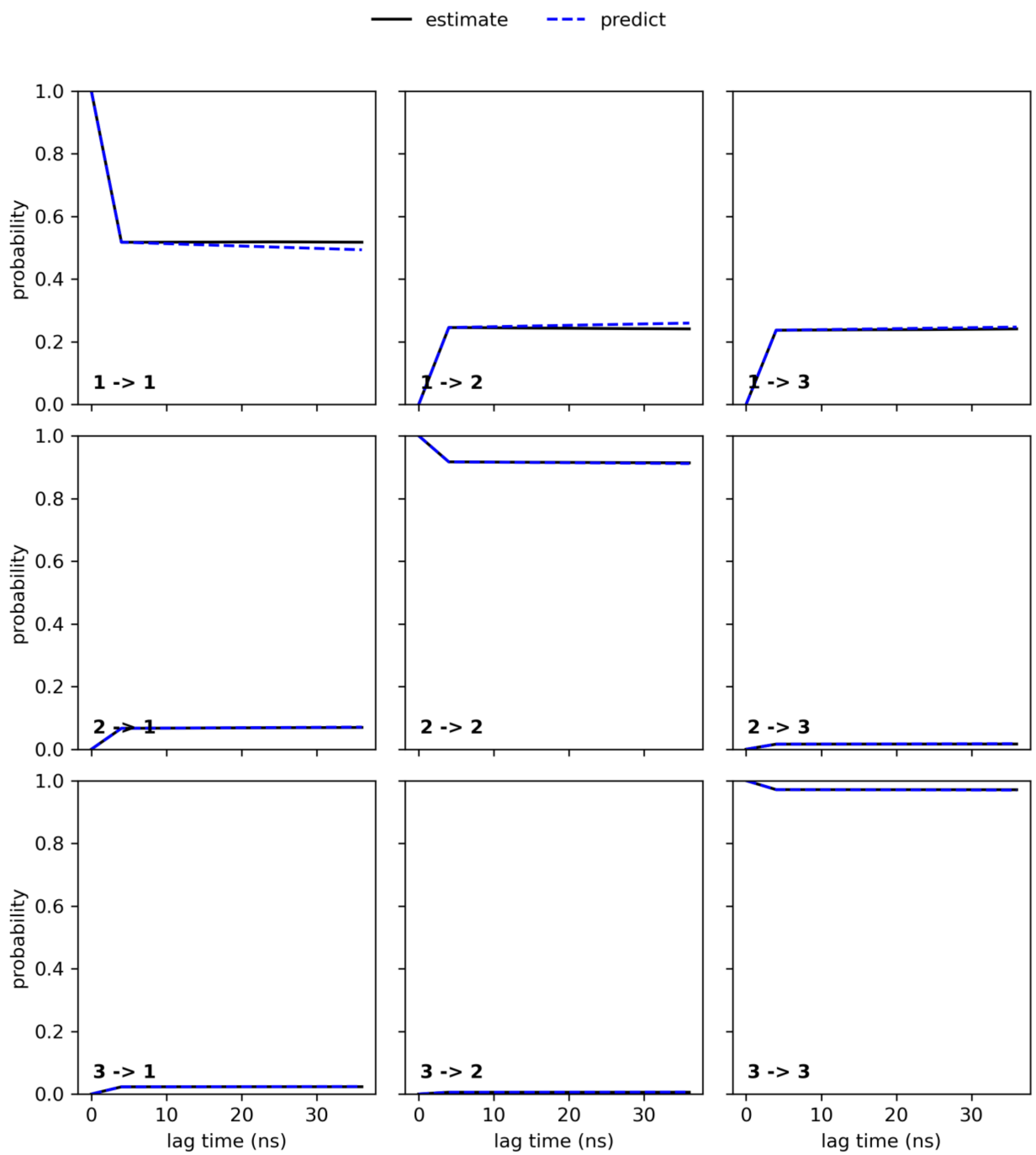


**Figure S10. Chapman–Kolmogorov test for the MSM of SoPIP2:PLPG conformational dynamics.**

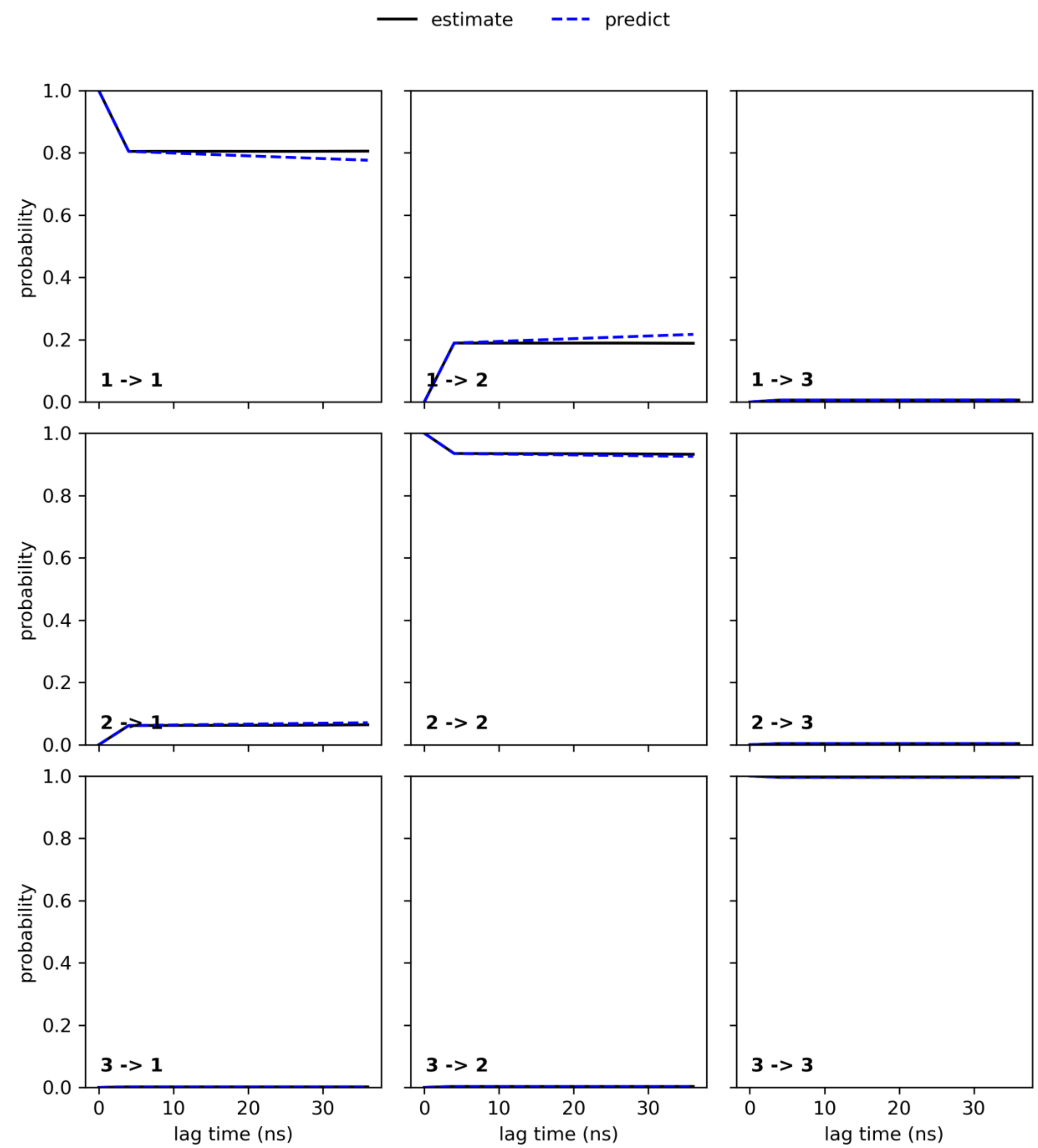


**Figure S11. Chapman–Kolmogorov test for the MSM of SoPIP2:LLPC conformational dynamics.**

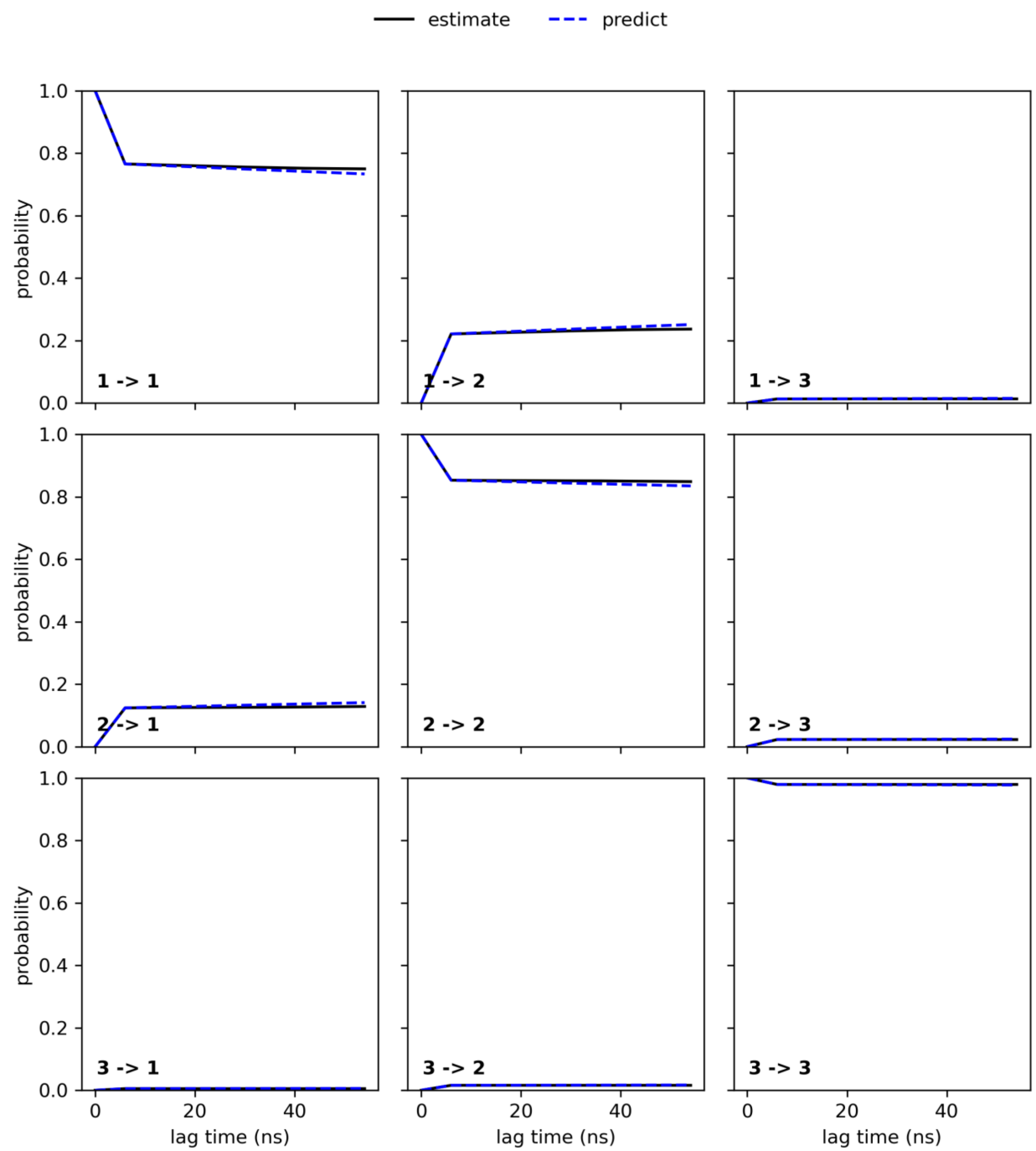


**Figure S12. Chapman–Kolmogorov test for the MSM of SoPIP2:LLPE conformational dynamics.**

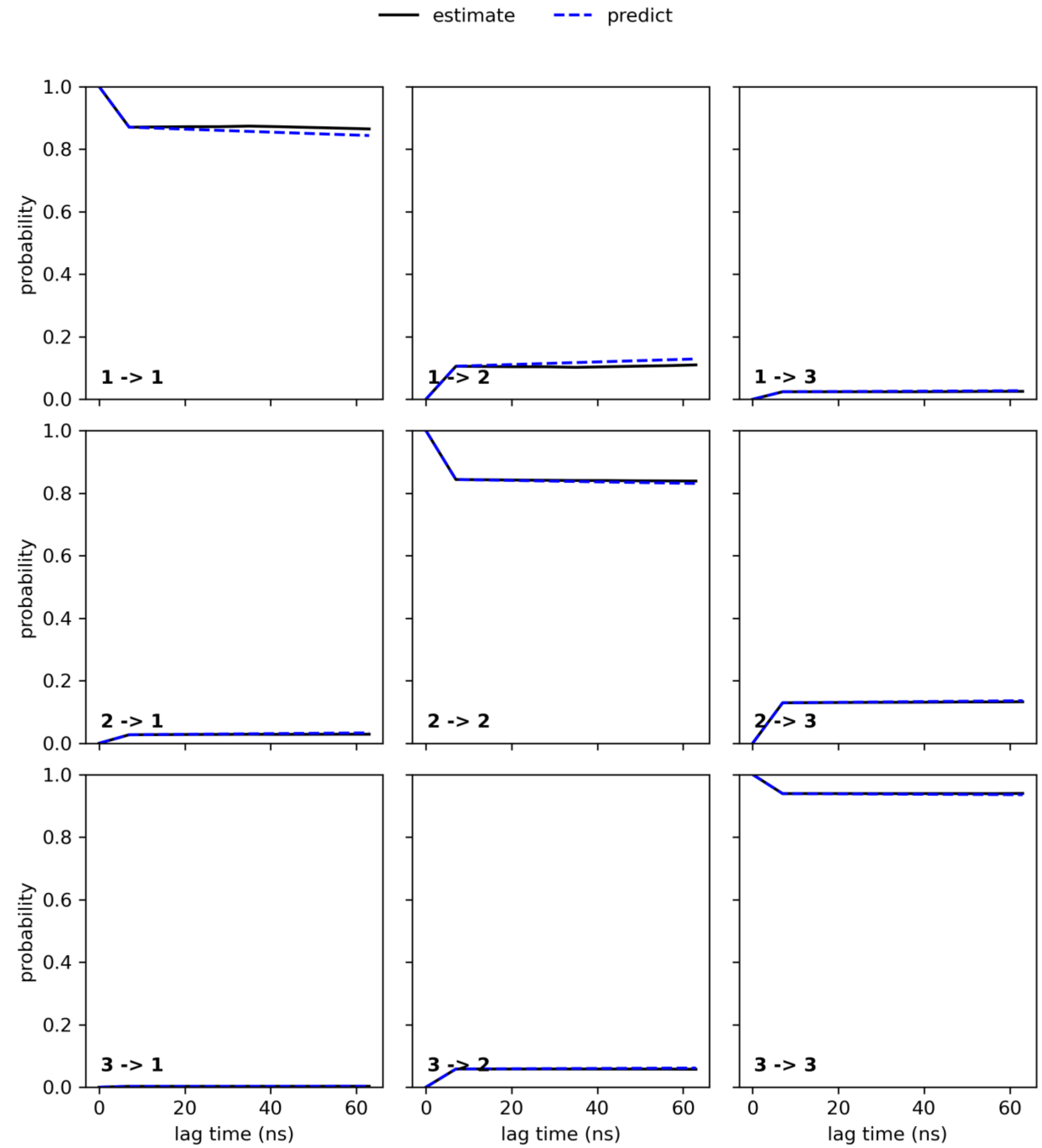


**Figure S13. Chapman–Kolmogorov test for the MSM of SoPIP2:LLPG conformational dynamics.**

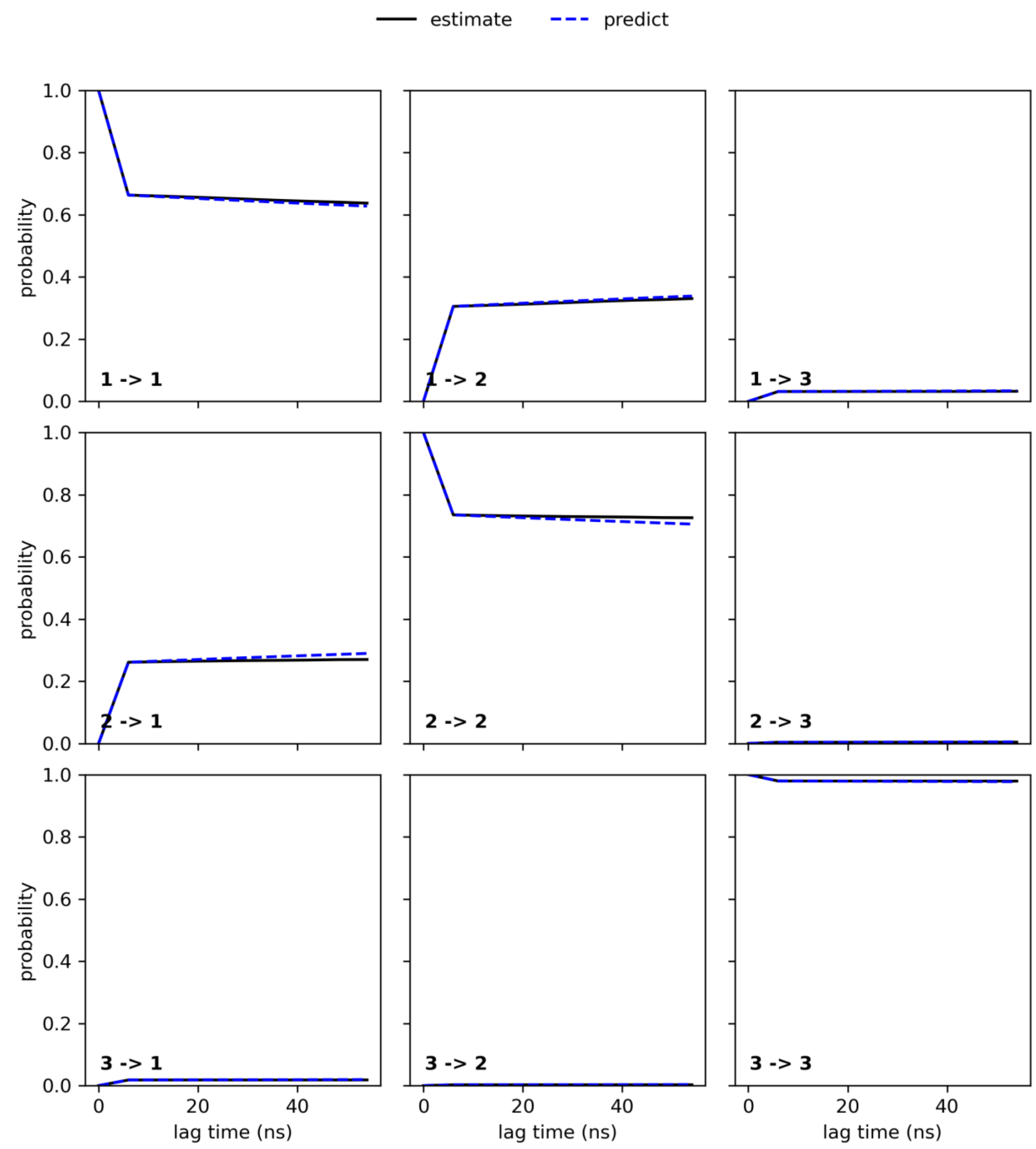


**Figure S14. Chapman–Kolmogorov test for the MSM of SoPIP2:complex conformational dynamics.**

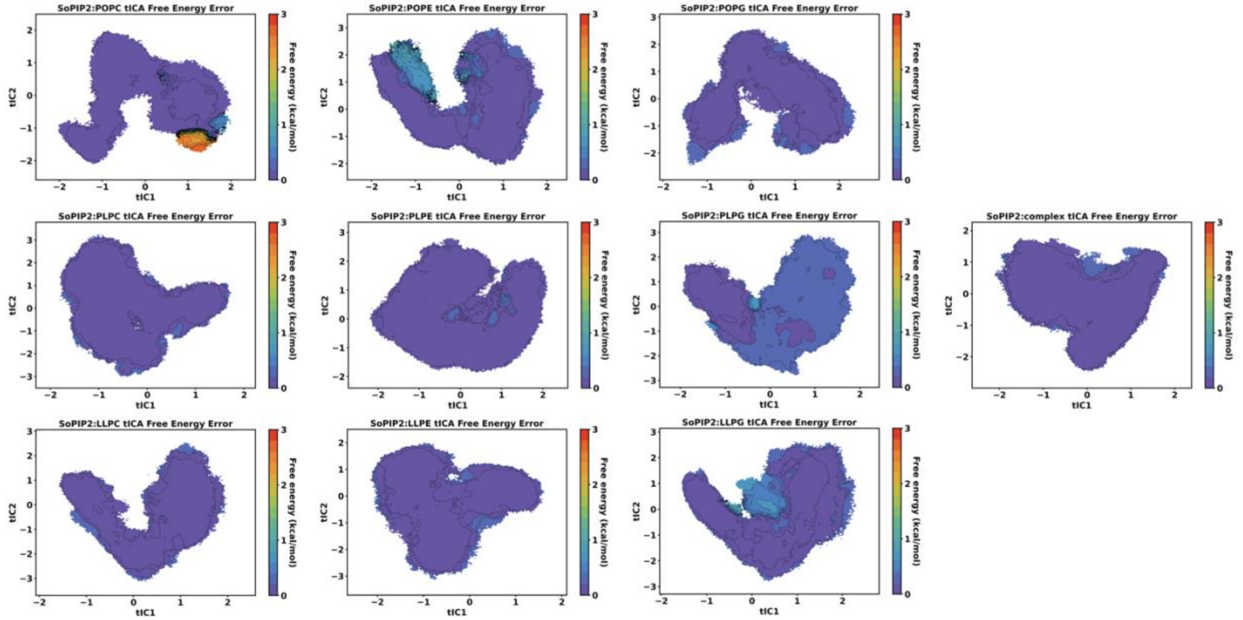


**Figure S15. Free energy error from the 200-sample bootstrapping protocol for each SoPIP2:bilayer system.**

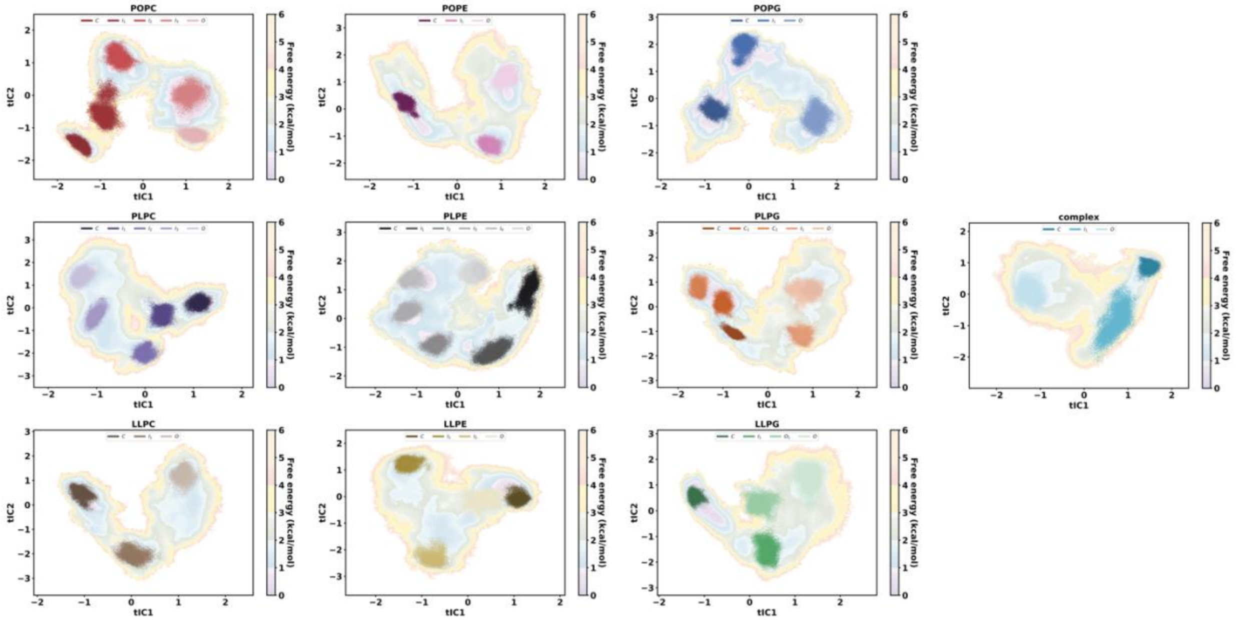


**Figure S16. Selected continuous trajectories for the water transport analysis based on tICA macrostate identification.**

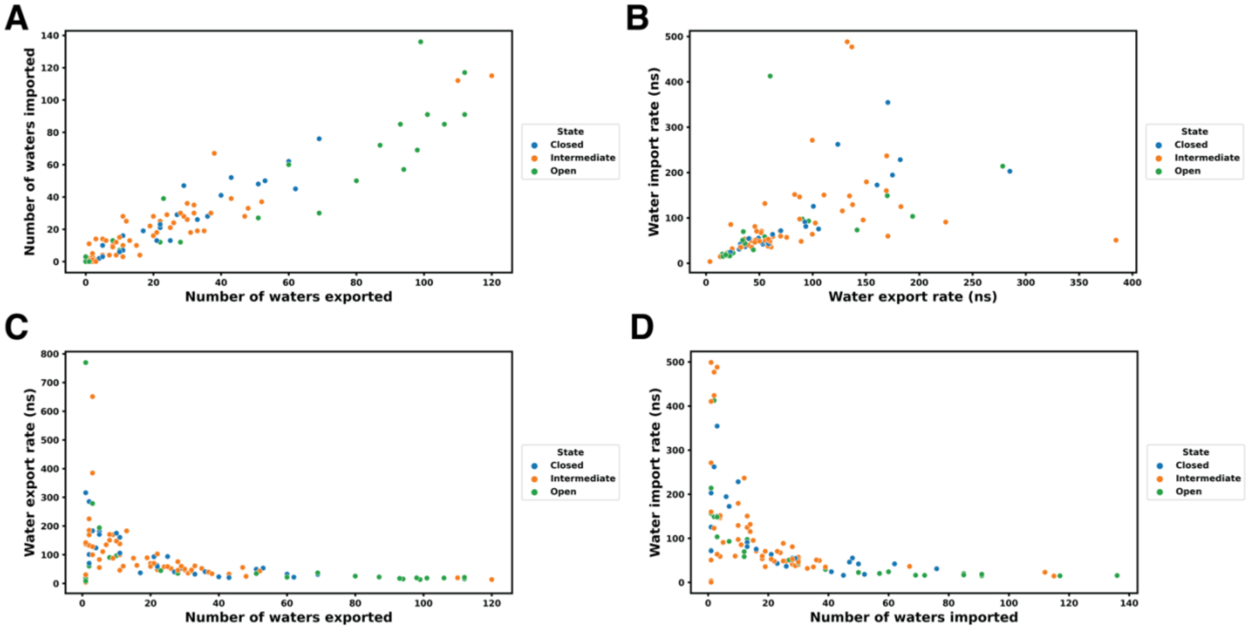


**Figure S17. Relationship between number of waters transported and the rate of transport.** (A) The positive correlation between the number of waters transported inside (imported into) the pore versus the number of waters transported outside (exported from) the pore. (B) Slight correlation between the export and import rate of transported waters. (C) Relationship between the rate and number of waters for export processes. (D) Relationship between the rate and number of waters for import processes.

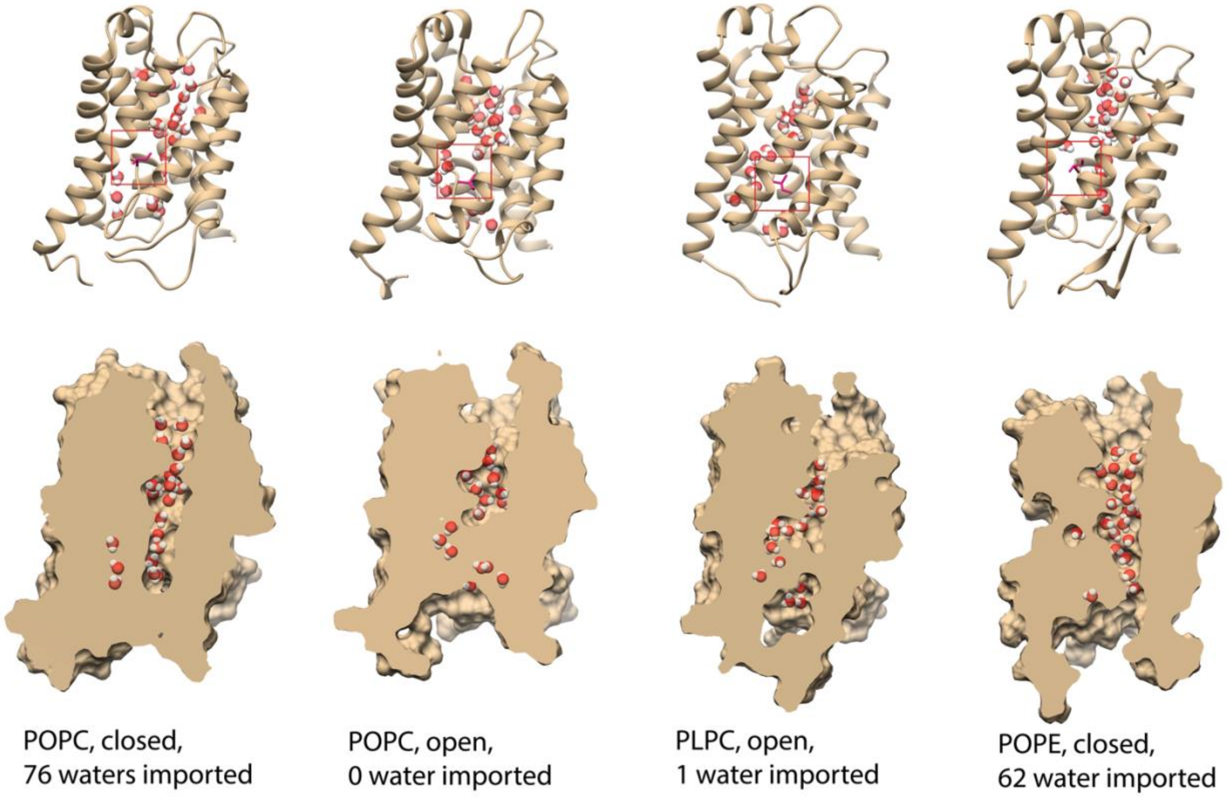


**Figure S18. Pore cavity structure with respect to loop D conformation throughout different types of example transport cases.**

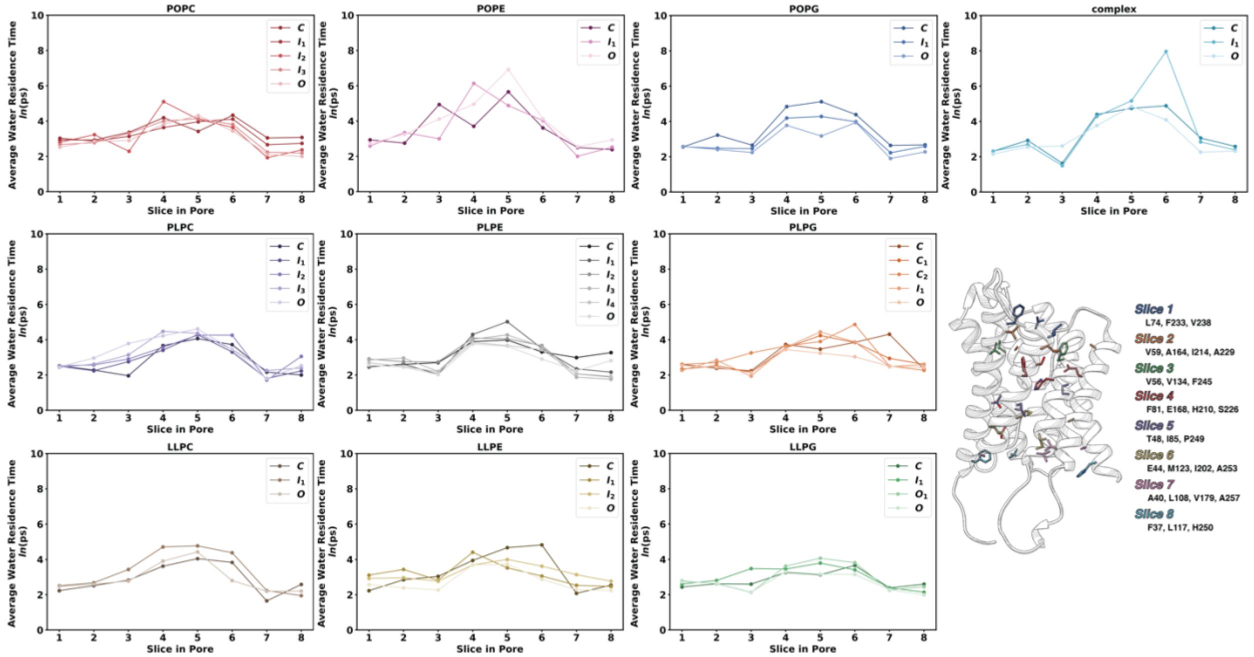


**Figure S19. Residence time of water molecules at each slice of the SoPIP2;1 pore.** The protein is separated into eight lateral slices along the vertical z-vector of the central pore. Each slice is defined as a cylinder whose center is given by the center of geometry from the $C_\alpha$ of 3-4 residues shown in the snapshot. The height of the cylinder is ± 2 Å from the cylinder center. The radius of the cylinder is 8 Å. The selection is done dynamically with MDAnalysis 2.0.0 "cyzone" geometric atom selection. The time for which each water molecule spends continuously in a given slice is averaged for all waters found in the slice throughout the trajectory to give the residence time. Most of the residence time landscape follows a similar pattern: water molecules (1) enter in a disordered manner and spend around ~9 ps in Slices 1-2; (2) spend less amount of time in Slice 3 (the selectivity region); then (3) remain in Slices 4-6 for a significantly higher time of ~65-80 ps (the NPA region and pore center); and finally (4) spend ~6 ps in the intracellular regions of the channel. Deviations from this pattern include all macrostates of SoPIP2:POPE and the intermediate state of SoPIP2:complex. Reasons for the intermediate state of SoPIP2 in the complex membrane could be a water molecule stuck in the opening of the second pathway. Meanwhile, the POPE bilayer induces a general difficulty in transport due to its high acyl chain order parameters and thickness (Figure S20-21).

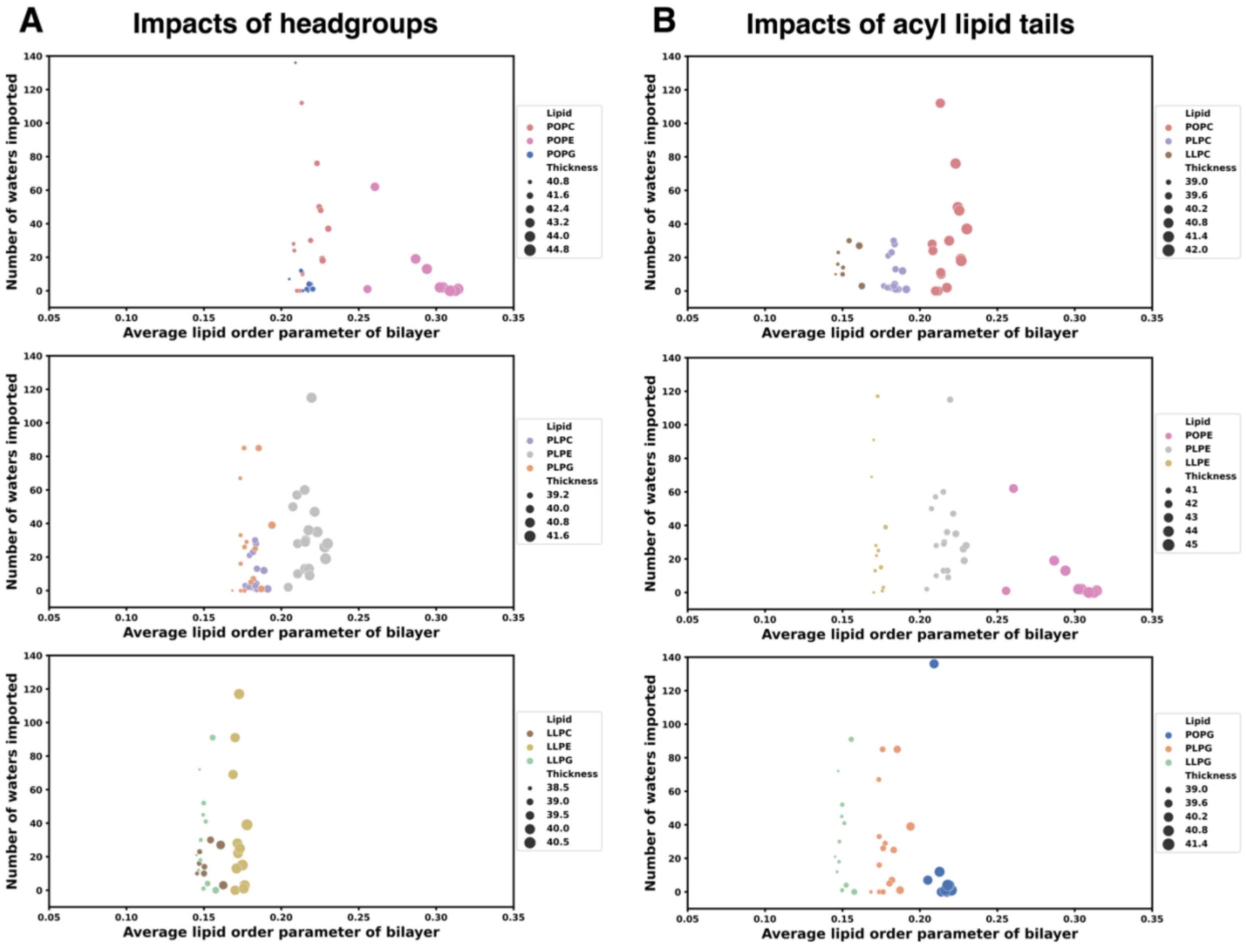


**Figure S20. Dissecting influence of lipid headgroups or acyl chains on SoPIP2;1 protein function.** Colors represent each homogeneous lipid bilayer system. Dot sizes correspond to the average thickness of the membrane bilayer in each 100-ns trajectory.

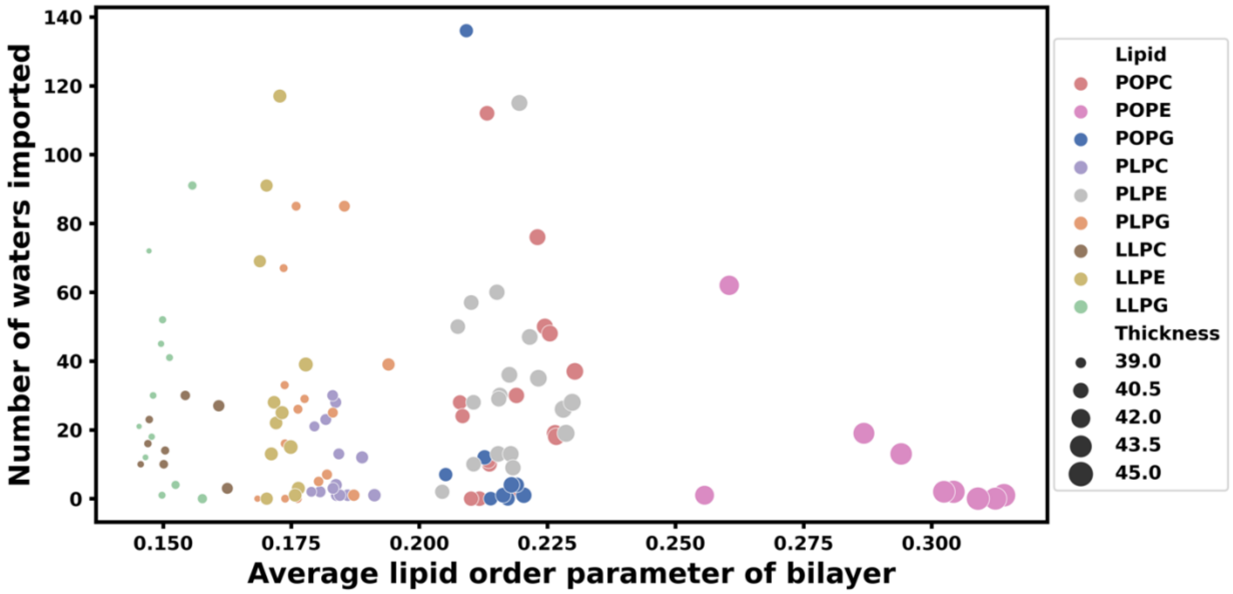


**Figure S21. Number of waters imported versus average lipid order parameter for all selected trajectories belonging to each SoPIP2:bilayer macrostate.** Colors represent each homogeneous lipid bilayer system. Dot sizes correspond to the average thickness of the membrane bilayer in each 100-ns trajectory.

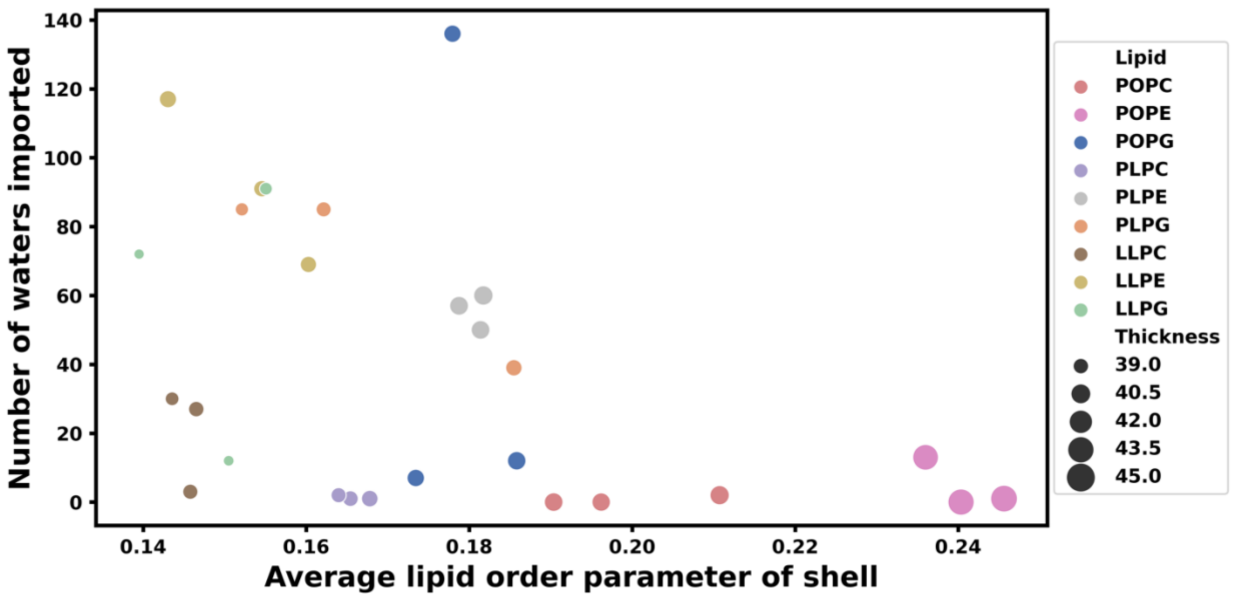


**Figure S22. Number of waters imported versus the average lipid order parameter of the annular shell lipids of the open states.**

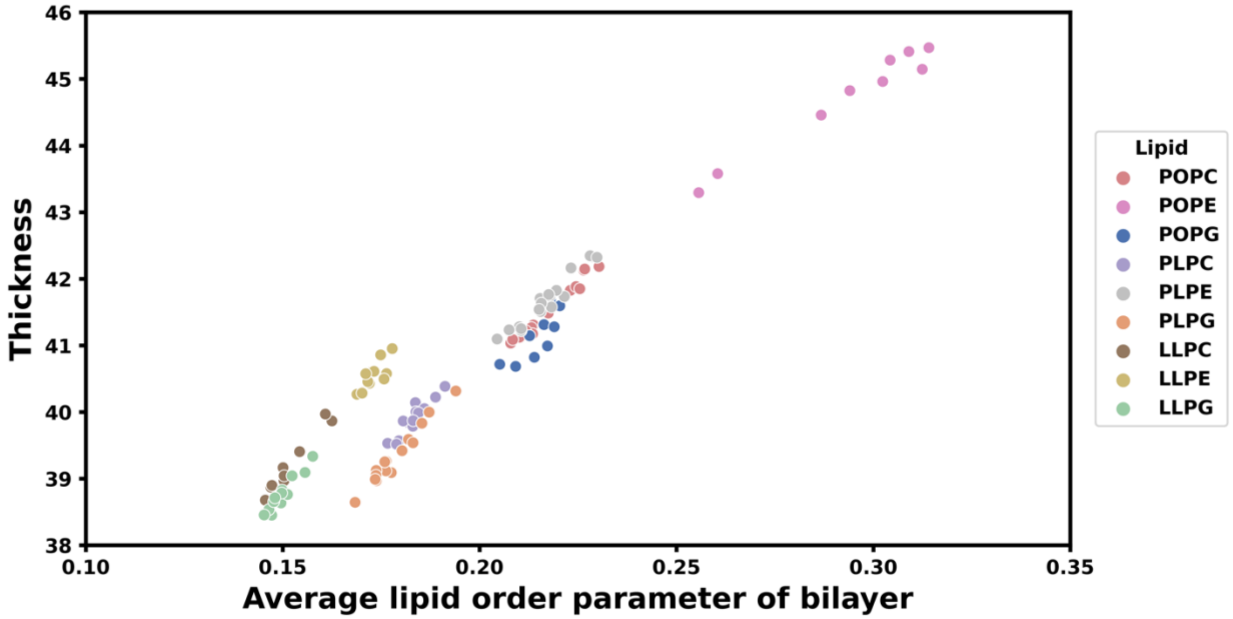


**Figure S23. Correlation between the thickness and order parameter of the lipids for each homogeneous SoPIP2:bilayer macrostate.**

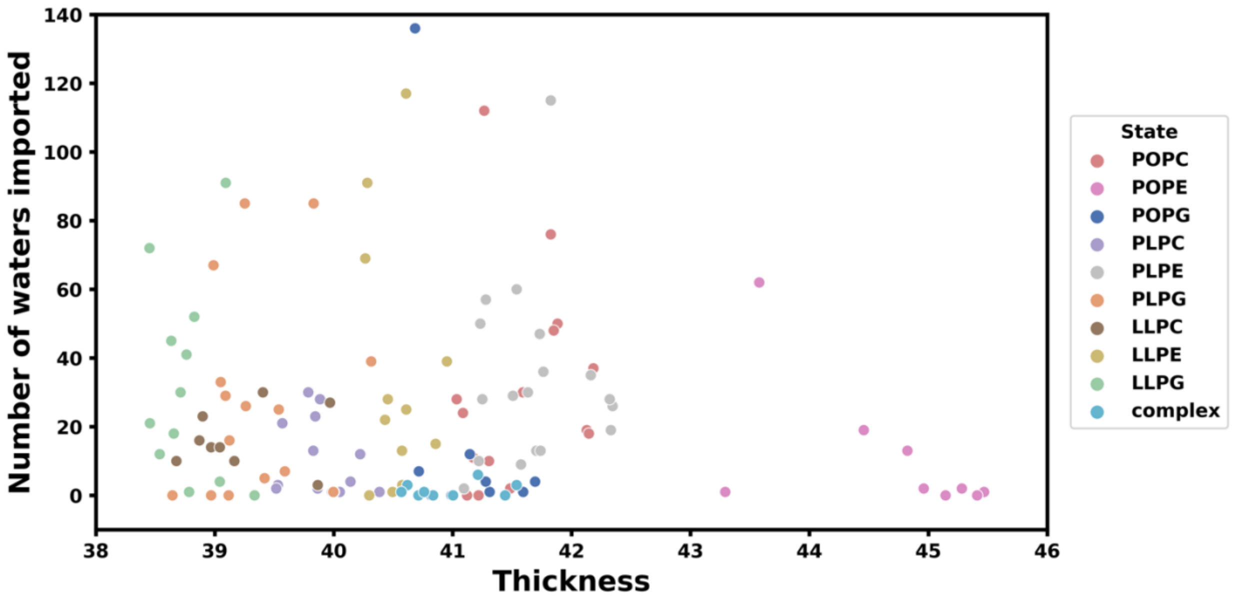


**Figure S24. Number of waters imported versus thickness for each SoPIP2:bilayer macrostate.**
